# Metagenomic classification of ancient viruses

**DOI:** 10.1101/2025.11.07.687203

**Authors:** Luís L. Marques, Armando J. Pinho, Diogo Pratas

**Affiliations:** IEETA/LASI - Institute of Electronics and Informatics Engineering of Aveiro, and University of Aveiro, 3810-193 Aveiro, Portugal; DETI - Department of Electronics, Telecommunications and Informatics, University of Aveiro, 3810-193 Aveiro, Portugal; DoV - Department of Virology, University of Helsinki, 00014 Helsinki, Finland

**Keywords:** Metagenomics, Metagenomic Classification, ancient DNA, data compression, ancient viruses

## Abstract

Ancient DNA (aDNA) sequences present unique challenges for taxonomic classification due to extreme fragmentation (reads 20-100 bp), end-biased cytosine deamination, and high contamination rates. Conventional metagenomic classifiers based on exact *k*-mer matching or alignment lose discriminative power on such short and damaged reads, limiting the analysis of paleogenomic samples. We present FALCON2, a compression-based metagenomic classifier that leverages position-aware finite-context models to maintain high accuracy on degraded viral ancient viruses. FALCON2 consolidates the capabilities of its predecessor, FALCON-meta, into a unified executable with enhanced features including model persistence, direct processing of compressed inputs, multiple file handling, and optional pre-filtering methodologies for contaminated samples. Under controlled benchmarking with database, taxonomy, and thread parity on simulated viral datasets, FALCON2 achieved an Area Under the Curve of Receiver Operating Characteristic (AUC-ROC) of 0.999, an Area Under Precision-Recall Curve (AUPRC) of 0.968, and an *F*_1_-score of 0.918, substantially outperforming Centrifuge (AUPRC = 0.625), Kraken2 (AUPRC = 0.184), and CLARK-S (AUPRC = 0.013) on pooled micro-averaged metrics. FALCON2’s advantage is most pronounced on ultra-short reads (20-40 bp), where exact *k*-mers become sparse. FALCON2 pre-filtering at threshold 0.7 improved precision by 10 percentage points with negligible recall loss. FALCON2 runs on systems with 4-8 GB RAM for typical analyses. FALCON2 is freely available at https://github.com/cobilab/FALCON2 under GPL v3 license.

## 1 Introduction

Ancient DNA (aDNA) recovered from archaeologic, paleontologic and historical specimens yields key insights into extinct species, ancient microbiomes and the evolutionary processes that shaped them [13, 3].

However, aDNA sequences are typically characterized by extreme degradation: fragments are generally 20–100 bp in length, exhibit elevated cytosine deamination rates (C → T and G → A substitutions at read termini) and are often contaminated by modern and environmental DNA [5, 1, 18].

These properties compromise the reliability of conventional metagenomic classifiers, which rely on exact *k*-mer matches or long alignment anchors. In current practice, taxonomic classification of shotgun metagenomes follows two main paradigms: (i) exact or reduced *k*-mer methods with Lowest Common Ancestor (LCA) post-processing (e.g., Kraken2, Centrifuge, CLARK) [20, 7, 9]; and (ii) alignment-based LCA pipelines, such as MALT used in conjunction with MEGAN, typically relying on FM-index/BWT aligners [4].

These pipelines are commonly paired with aDNA-aware mapping and processing frameworks that mitigate short, damaged reads [17, 16]. These approaches perform strongly on long, low-error reads but lose discriminative power as fragments shorten and post-mortem damage increases, precisely the regime characteristic of aDNA, because exact *k*-mers and long alignment anchors become sparse.

Marker-gene workflows (e.g., 16S/18S/ITS) and downstream functional imputation are powerful but target specific loci, making them less suitable for shotgun aDNA where fragmentation and damage are pervasive [2, 8].

Compression-based methods, including FALCON-meta, avoid the need for long exact substrings by comparing relative compressibility between reads and references, retaining signal on ultra-short, damaged fragments [12].

FALCON-meta [12] introduced a compression-based approach to taxonomic classification, using Finite-Context Models (FCMs) and Normalized Relative Similarity (NRS) to quantify sequence relationships without requiring exact substring matches. This approach demonstrated robustness to short and damaged reads, but was implemented as a fragmented codebase of multiple scripts, lacked model reuse capabilities, and did not address contamination filtering.

We present FALCON2, the successor to FALCON-meta, engineered as a unified, production-ready tool for ancient viral metagenomics. FALCON2 incorporates finite-context models, model persistence via serialization, native handling of compressed inputs (FASTA/FASTQ, gzip), multiple file processing, and optional integration of a lightweight compression-based pre-filter for removing contaminant reads. Benchmarking on synthetic viral datasets with factorial combinations of read length (20-100 bp), deamination rate (0-30%), and sequencing depth (1-60*×* ) shows that FALCON2 outperforms established nucleotide-based classifiers on short and damaged fragments while maintaining practical computational requirements.

## 2 Methods

FALCON2 classifies metagenomic reads by computing NRS scores between query sequences and reference genomes [12]. For each query sequence *x*, FALCON2 trains multiple FCMs on a reference sequence *y*, then encodes *x* using these reference-derived (frozen) models to obtain the compressed length *C*(*x*||*y*).

The method uses a cooperative mixture of multi-order FCMs and substitution-tolerant Markov models, which estimate symbol probabilities based on preceding context and allow for a degree of mismatch tolerance. The efficiency of relative compression depends on how well the reference-trained model collects and organizes information so that questions about the target can be answered with as few bits as possible.

This relative compression setup implies that the compressor cannot exploit intra-target redundancies, only information learned from the reference can reduce code length. Intuitively, if the reference describes the target well, the required relative information is small; if the target is not related, the encoder approaches the maximum number of bits.

Formally, the Relative Similarity (RS) is defined as

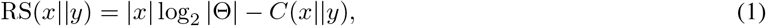

and the normalized version as

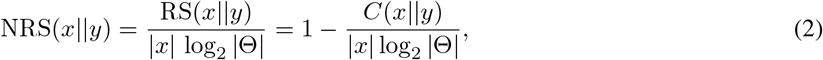

where |*x*| is the size of the sequence and Θ is the size of the alphabet (4 for DNA). In read-level classification, FALCON2 assigns *x* to the reference from the database that produces the highest NRS. Additional details can be found at Supplementary Section 1.

For composition profiling (the `meta`subcommand), we instead treat the sample as a bag/concatenation of reads *X* and compute NRS(*y*_*i*_ || *X*) for each reference *y*_*i*_ in the database (*y*_1_, *y*_2_, …), thus quantifying how well the sample explains each reference and allowing global composition estimates [12, 11, 10]. For efficiency, the highest-scoring references (top-*K* by NRS) are cached and reused across computations.

### 2.1 Unified executable

FALCON-meta exposed functionality through several binaries: for global composition (FALCON), local-similarity filtering and visualization (FALCON-filter, FALCON-filter-visual), and inter-reference similarity (FALCON-inter, FALCON-inter-visual) [12, 11, 10]. FALCON2 consolidates these into a single production-ready executable that presents equivalent capabilities as subcommands: `meta`for composition, `filter`/`fvisual`for segmentation and visualization of local profiles, and `inter`/`ivisual`for computing and rendering genome–genome similarity. The interface adds practical conventions, native streaming of gzip-compressed FASTA/ FASTQ and colon-separated multi-file tokens for paired reads or multi-FASTA references, as well as model persistence and stricter reproducibility guaranties.

In routine use, composition runs as `FALCON2 meta [options] READS_GROUP DB_GROUP`, where each positional argument is a single colon-separated token (e.g., `R1.fq.gz:R2.fq.gz`for paired-end reads and `ref1.fa:ref2.fa`for multiple references). Inputs may be gzip compressed and are streamed directly, avoiding prior decompression. When local profiling is required, `meta`emits a profile in-process (e.g., with `-Z -y profile.tsv`), which is then segmented by `filter`and rendered by `fvisual`without format conversion. For database inspection and quality control, `inter`computes a genome-by-genome similarity matrix from the same multi-FASTA reference set, and `ivisual`produces a publication-ready heat map. Per-command help follows standard conventions and is available via `FALCON2 <command> -h`.

FALCON2 operates directly on data from sequencers, independently of coverage, and accepts both assembled references and non-assembled read sets. Although FALCON2 is designed for ancient viral metagenomics, it scales to large organismal databases (viral, bacterial, archaeal, fungal) as well as custom collections.

### 2.2 Model persistence and reuse

Trained FCMs can be serialized to `.fcm`files and reloaded in subsequent runs, enabling computational reuse in multi-sample analyses or when the reference database remains constant. Model persistence is implemented via the `-S`(save) and `-L`(load) flags, combined with `-M`to specify the model path. This functionality reduces steady-state inference time and ensures reproducibility across runs. For reproducibility, `-T`enables train-only runs and `-I`prints model metadata (tool version, reference snapshot hash, key parameters). Reloading (`-L -M`) enforces basic compatibility checks so that stale or mismatched models are rejected.

### 2.3 Pre-filtering

FALCON2 optionally integrates a compression-based pre-filter for contaminated samples. It computes approximate similarity scores between reads and a contaminant library (e.g., *E. coli*, human), retaining only reads with similarity below a configurable threshold *τ* [19]. Reads exceeding *τ* are excluded before FALCON2 classification, reducing computational load and improving precision. The pre-filter is activated with `-mg`, with threshold controlled by `-mt`(default 0.9; recommended 0.6–0.7 for aDNA).

### 2.4 Output and parallelization

Outputs are tabular files reporting NRS scores and taxonomic assignments for each read. Multi-threading is controlled via the `-n`parameter (default: all available cores). Internally, FALCON2 uses cache-aware hashing to memoize local probabilities and maintains a top-*K* cache of the highest NRS values across passes for speed. The compression depth parameter `-l`controls model fidelity; `-l 47`was used in all benchmarking to maximize discrimination on short reads. The tool is freely available, under the GPLv3 license, at https://github.com/cobilab/FALCON2.

## 3 Benchmark

### 3.1 Experimental design

We benchmarked FALCON2 against Centrifuge [7], Kraken2 [20] and CLARK [9] under strict parity conditions (details in Supplementary Section 2). All tools used identical reference databases (NCBI RefSeq-Viruses plus *E. coli* K-12 and human mitochondrial DNA as contaminants), as well as the same NCBI taxonomy snapshot, and fixed thread counts (n=8). Synthetic datasets were generated via Gargammel [14] (for aDNA fragmentation and deamination) and ART [6] (for sequencing errors), spanning combinations of read length (20, 40, 60, 80, 100 bp), deamination rate (0.0, 0.1, 0.2, 0.3), and sequencing depth (1, 5, 10, 20, 40, 60 *×* ). Ground truth was extracted from simulation metadata, and species-level precision, recall, *F*_1_-score, AUPRC, and AUC-ROC were computed. AUPRC was prioritized over AUC-ROC due to class imbalance [15].

### 3.2 Results

Table 1 summarizes pooled micro-averaged performance across all experimental conditions. Details are in Supplementary Section 3-9. FALCON2 achieved the highest AUPRC (0.968), *F*_1_-score (0.918), and AUC-ROC (0.999), substantially outperforming Centrifuge (AUPRC = 0.625, *F*_1_ = 0.738), Kraken2 (AUPRC = 0.184, *F*_1_ = 0.372), and CLARK (AUPRC = 0.013, *F*_1_ = 0.103). The advantage was most pronounced at read length 20 bp, where FALCON2’s AUPRC exceeded Centrifuge by 0.34 and Kraken2 by 0.75 (Figure 1). At read length 100 bp with low deamination (0.0), performance differences narrowed (ΔAUPRC ≈ 0.06), consistent with the hypothesis that *k*-mer methods reassert efficiency on long, intact reads.

**Table 1:** Pooled micro-averaged performance metrics across all experimental conditions. Best values in bold. Time (Wall) is in minutes and RAM (peak) in in GigaBytes.

| Tool | Precision | $F_1$ | AUCROC | AUPRC | Time | RAM |
| --- | --- | --- | --- | --- | --- | --- |
| **FALCON2** | **0.932** | **0.918** | **0.999** | **0.968** | 0.88 | 1.90 |
| Centrifuge (default) | 0.753 | 0.738 | 0.914 | 0.625 | 0.20 | **0.07** |
| Kraken2 (default) | 0.508 | 0.372 | 0.666 | 0.184 | **0.04** | 0.45 |
| CLARK (runtime via CLARK-S) | 0.116 | 0.103 | 0.548 | 0.013 | 0.20 | 1.57 |

**Figure 1.**
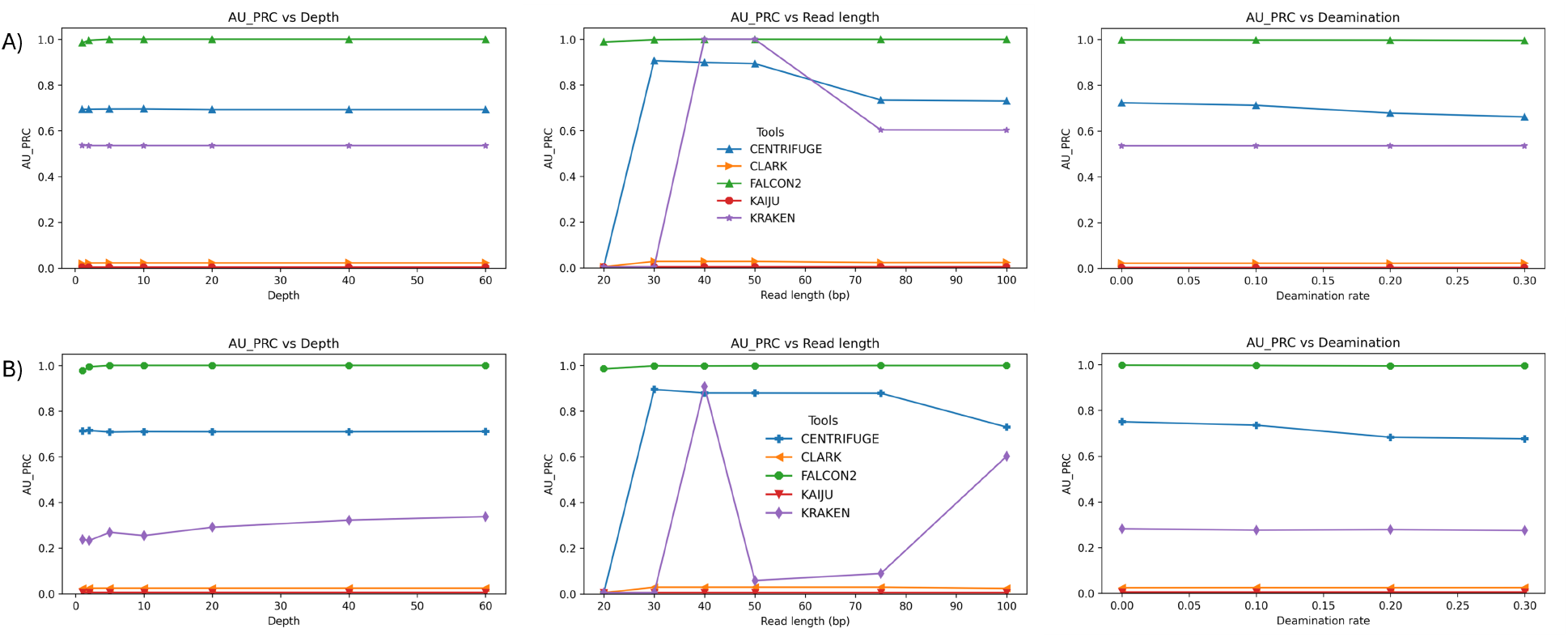
AUPRC across depth, read length, and deamination with contamination applied. A) AUPRC before trimming and B) AUPRC after trimming.

Pre-filtering at threshold 0.7 increased precision from 0.85 to 0.95 while recall declined minimally from 0.90 to 0.87. The retained fraction *k*(*τ* ) decreased from 0.70 to 0.30, indicating 70% of reads were filtered. An equivalence test confirmed that disabling filtering (*τ* = 1.0) produced byte-identical outputs, validating orchestration integrity.

In summary, FALCON2’s compression-based approach maintains discriminative capacity on short and damaged aDNA reads where exact *k*-mer methods degrade. The integration of position-aware models, model persistence, and pre-filtering provides a robust, production-ready tool for ancient DNA metagenomics. The benchmarking framework enforces strict parity conditions and is fully reproducible via the archived scripts and data (https://doi.org/10.5281/zenodo.17291215).

### 3.3 Computational resources

FALCON2 exhibited higher runtime (median 0.88 min per sample) and memory usage (median 1.90 GB) than Kraken2 (0.04 min, 0.45 GB) and Centrifuge (0.20 min, 0.07 GB), reflecting the computational cost of FCM. However, absolute times remain practical for typical metagenomic workflows. On-disk footprint for the viral reference database was 75 MB (FALCON2), 45 MB (Centrifuge), and 255 MB (Kraken2). When using model persistence, build costs shift to a one-time training phase, and steady-state inference accelerates.

## 4 Conclusions

FALCON2 advances ancient viral metagenomic classification with compression-based models robust to fragmentation and deamination. Under controlled benchmarking, FALCON2 achieved superior AUPRC, *F*_1_, and AUC-ROC compared to established classifiers, with the largest advantages on ultra-short reads (20-40 bp). The tool’s unified architecture, model reuse, and contamination filtering capabilities establish FALCON2 as an open-source solution for ancient viral metagenomic analysis.

