## Supplementary Material for "Metagenomic classification of ancient viruses"

#### 1 Normalized Relative Similarity (NRS)

We quantify the information content of a sequence and compare two sequences through a relative, compression-based lens grounded in Kolmogorov complexity. Following [1], effective computation is formalized by Turing machines. We fix a universal machine (left implicit in the notation) and use the convention that $p = x$ means "program $p$ prints $x$ and halts," $p(y) = x$ means "$p$, given $y$ as auxiliary data, prints $x$ and halts," and $p = (x, y)$ means "$p$ prints the pair $(x, y)$ and halts." With this convention, for a finite string $x$ the Kolmogorov complexity is

$$K(x) \;=\; \min_{p=x} \ell(p).$$

The conditional Kolmogorov complexity of $x$ given $y$ is

$$K(x \mid y) \;=\; \min_{p(y)=x} \ell(p),$$

and the joint complexity is

$$K(x, y) \;=\; \min_{p=(x,y)} \ell(p).$$

These definitions realize Kolmogorov's proposal to measure information via the size of the most concise effective description [2], building on Solomonoff's inductive framework [3, 4] and Chaitin's program-size formalization [5]; see [6] for a comprehensive treatment. Whereas Shannon's entropy quantifies average information under a source distribution [7, 8], Kolmogorov complexity assigns information to individual objects. Standard relationships (customarily stated up to lower-order additive terms) include the chain rule

$$K(x, y) \;=\; K(x) \;+\; K(y \mid x)\,.$$

Since the exact complexities above are not computable in general, we employ compression-based proxies. To compare sequences with $y$ used strictly as side information for $x$, we adopt a frozen-model viewpoint: a compressor first learns a model from $y$ within finite time, freezes it, and then encodes $x$ *using only that frozen model.* Writing $C(x \,\|\, y)$ for this relative compressed length aligns the approach with relative-information ideas for individual sequences [9] and with compression-based similarity and distance measures such as information distance and its normalized variants [10, 11, 12]. Let $\Theta$ be the alphabet and $|\Theta|$ its size, and let $|x|$ be the number of symbols in $x$. The symbol-wise uninformed baseline is $|x| \log_2 |\Theta|$ bits; if $y$ carries no useful structure for $x$, a competent frozen-model compressor will produce $C(x \,\|\, y)$ close to this baseline, whereas if $y$ captures regularities that also govern $x$, the code will be substantially shorter. In this setting, the Relative Similarity (RS) is

$$\mathrm{RS}(x \,\|\, y) \;=\; |x| \, \log_2 \bigl|\Theta\bigr| \;-\; C(x \,\|\, y), \tag{1}$$

and the normalized version is

$$\mathrm{NRS}(x \,\|\, y) \;=\; \frac{\mathrm{RS}(x \,\|\, y)}{|x| \, \log_2 \bigl|\Theta\bigr|} \;=\; 1 \;-\; \frac{C(x \,\|\, y)}{|x| \, \log_2 \bigl|\Theta\bigr|}. \tag{2}$$

When $x = y$ over the same alphabet and the model class is expressive enough, the frozen model encodes $x$ at very low extra cost and $\mathrm{NRS}(x \,\|\, y)$ is close to one; when $x$ is algorithmically unrelated to $y$, the relative code length approaches the baseline and $\mathrm{NRS}(x \,\|\, y)$ is close to zero. For a concrete illustration over a binary alphabet, if $y$ is long enough for the compressor to learn a Bernoulli bias $p$ and $x$ is independently drawn from the same source, then the relative code length is about $|x| \, H_2(p)$, yielding $\mathrm{NRS}(x \,\|\, y) \approx 1 - H_2(p)$, consistent with probabilistic coding theorems while remaining grounded in the objectwise framework of algorithmic information theory [7, 8, 2, 5, 6].

### 2 Benchmarking Setup

In the following sections, we describe the architecture, how to build the database, cleaning and trimming, parameters and computing environment for the inference of metagenomic composition. Regarding the default aligning parameters, quantifying somewhat dissimilar sequences by alignment methods is problematic, due to the need of fine-tuned thresholds, considering relaxed edit distances and consequent need of very high computational resources.

#### 2.1 Architecture

The benchmarking pipeline (Figure S1) follows a structured sequence to ensure fair and reproducible evaluation. First, we construct a standardized reference database from an NCBI RefSeq snapshot (comprising viral, bacterial, archaeal, and fungal sequences), consolidate it into a single multi-FASTA file, and map accessions to their corresponding NCBI Taxonomy identifiers. All classifiers use this common reference to eliminate variability due to database content or update timing.

Second, we generate synthetic reads with known ground truth from this reference using Gargammel (to model ancient DNA fragmentation and deamination) and ART (to model contemporary sequencing error), thereby covering both aDNA and modern metagenomic regimes.

Third, each classifier is executed against the same reference/taxonomy snapshot under identical operating conditions (threads and reporting settings), producing taxonomic assignments and relative-abundance profiles. Finally, predictions are compared to the known ground truth using precision, recall, and $F_1$, and we record runtime and peak memory to assess computational efficiency alongside accuracy.

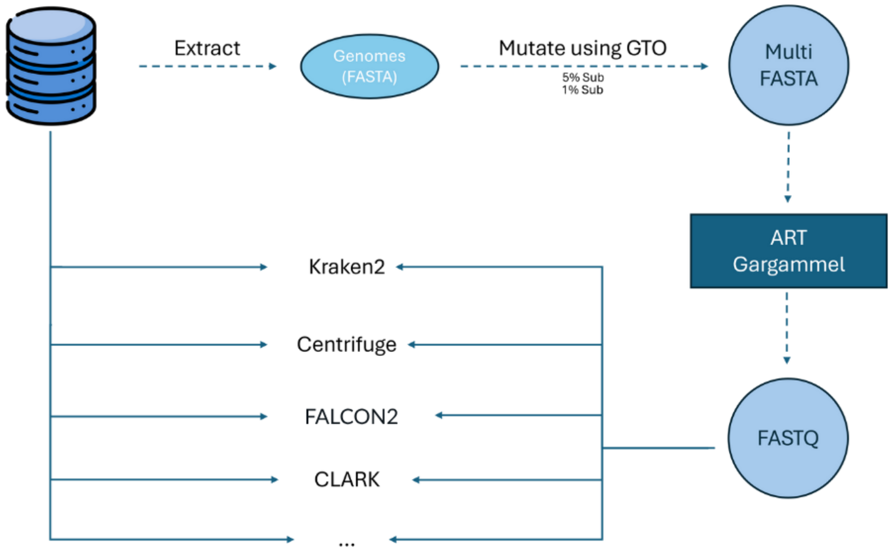


Figure S1: Representation of the Benchmarking Process. Genomes (FASTA) are extracted, mutated, simulated into reads with ART/Gargammel (FASTQ), and then benchmarked across classifiers (Kraken2, Centrifuge, FALCON2, CLARK).

### 3 Software and Environment

##### Hardware/OS

- **CPU:** `13th Gen Intel(R) Core(TM) i7-13700KF`, cores/threads: `12/12`.
- **RAM:** `19 GiB`.
- **OS/Kernel:** `Ubuntu 24.04.2 LTS`, kernel `6.8.0-85-generic`.
- **Storage:** `1 TB`.

##### Compilers/Interpreters

- **C/C++:** `gcc (Ubuntu 13.3.0-6ubuntu2~24.04) 13.3.0`; `GNU Make 4.3`.
- **Python:** `Python 3.12.3`.

**Profiling & provenance**

- **Runtime/memory:** `/usr/bin/time -v` for wall-clock, CPU time, peak RSS.
- **Hashing:** `/usr/bin/sha256sum` for binary/script checksums.

  Benchmark runs used `n = 8` threads.

**Other tools (benchmarks)** Executed from the `benchmark` conda environment.

- **Kraken2:** `2.1.3` (from `kraken2 -v`).
- **Centrifuge:** `1.0.4` (from `centrifuge --version`).
- **CLARK/CLARK-S:** `1.3.0` (from `CLARK -- help | head -n 1`)

**Dependencies**

- Entrez Direct

```
sh -c "$(curl -fsSL https://ftp.ncbi.nlm.nih.gov/entrez/entrezdirect/install-edirect.sh)"
```

- Gargammel

See: `https://github.com/grenaud/gargammel`

### 4 Database Setup

Download the reference genomes; you can use others and the following commands that mirror the process used for benchmarking.

#### 4.1 Download Reference Genomes

```
# Download viral genomes from NCBI
./download_references_ncbi.sh viruses
```

#### 4.2 Accession to TaxID Mapping

```
# From a file
./accession_to_taxid_v3.sh -f accessions.txt -e -o results.tsv

# Parse FASTA files in a directory
./accession_to_taxid_v3.sh -d /path/to/fastas -e -o taxids.tsv

# From comma-separated list
./accession_to_taxid_v3.sh -a "NC_001925.1,NC_000913.3" -e
```

#### 4.3 FALCON2

```
# Concatenate all reference sequences into a single FASTA file
./generate_reference_sequences.sh /path/to/reference_fastas input-sequences.fna

# Alternative: process compressed files (.gz) and concatenate
./process_gz_files.sh /path/to/reference_fastas input-sequences.fna

# Alternative: manual concatenation from decompressed files
cat /path/to/reference_fastas/*.fna > input-sequences.fna
```

##### 4.4 Kraken2

```
# Step 1: Convert FASTA headers to Kraken2 format (using assembly metadata) and save in
    kraken2_ready
./fasta_to_kraken2.sh viruses/viruses_sorted.tsv viruses/reference_fastas/

# Step 2: Download NCBI taxonomy
k2 download-taxonomy --db kraken2_db

# Step 3: Add files to library (batch process)
./kraken_add_to_library.sh kraken2_db kraken2_ready

# Alternative: add a single file
k2 add-to-library --file kraken2_ready/GCF_000837145.fna --db kraken2_db

# Step 4: Build the database
kraken2-build --build --db kraken2_db --threads 8
```

**Note:** Do *not* use `kraken2-build --add-to-library` (known issues). Prefer `k2 add-to-library`.

##### 4.5 Centrifuge

```
# Step 1: Create seqid2taxid conversion table
./fasta_to_seq2taxid.sh viruses/viruses_sorted.tsv viruses/reference_fastas/

# Step 2: Concatenate all sequences into single file
# Option A: from compressed files
./process_gz_files.sh viruses/reference_fastas/ input-sequences.fna
# Option B: from decompressed files
cat library/*/*.fna > input-sequences.fna

# Download NCBI taxonomy (if not already present from Kraken2 setup)
centrifuge-download -o taxonomy taxonomy

# Step 3: Build Centrifuge index
centrifuge-build -p 8 \
    --conversion-table seqid2taxid.map \
    --taxonomy-tree kraken2_db/taxonomy/nodes.dmp \
    --name-table kraken2_db/taxonomy/names.dmp \
    input-sequences.fna c_viruses
```

##### 4.6 CLARK-S

```
# Step 1: Create database directory structure
mkdir clark_viruses
mkdir clark_viruses/Custom

# Step 2: Decompress reference FASTA files (CLARK doesn't accept compressed files)
./decompress_fna_files.sh viruses/reference_fastas/ \
    viruses/decompressed_reference_fasta

# Step 3: Copy sequences to Custom folder
cp viruses/decompressed_reference_fasta/* clark_viruses/Custom/
# OR use rsync
sudo rsync -vahP viruses/decompressed_reference_fasta/ \
    CLARKV1.3.0.0/clark_viruses/Custom/

# Step 4: Define targets
./set_targets.sh clark_viruses custom

# Note: Build essentials required (if not already installed)
# sudo apt update && sudo apt install build-essential
```

#### 5 Simulate Ancient DNA Reads with Gargammel

Prepare reference data:

- **Endogenous DNA** (`endo/`): target organisms (e.g., concatenated viral genomes)
- **Bacterial contamination** (`bact/`): e.g., *E. coli* (GCF_000005845.2)

- **Human contamination** (`cont/`): human mitochondrial DNA (NC_012920.1)

```
# Run multiple parameter combinations
./run_multiple_simulations.sh

# Customizable parameters in script:
# - depths: (1 2 5 10 20 40 60)
# - read_sizes: (20 30 40 50 75 100 150)
# - deaminations: (0 0.1 0.2 0.3)
```

**Directory structure for Gargammel**

```
data/
|-- endo/   # Target DNA (viruses, etc.)
|-- bact/   # Bacterial contamination
`-- cont/   # Human contamination
```

### 6 Read Processing

```
./trim_fastq_files.sh input_dir output_dir
```

### 7 Run Classification Tools

Using the provided script utils, if using the simulated reads created by Gargammel.

#### 7.1 FALCON2

```
./falcon2_run.sh simulated_reads
```

#### 7.2 Kraken2

```
./kraken2_run.sh simulated_reads
```

#### 7.3 Centrifuge

```
./centrifuge_run.sh simulated_reads
```

#### 7.4 CLARK

```
# Classify samples
./clark_classify.sh simulated_reads output/CLARK

# Generate reports
./clark_run.sh output/CLARK
```

### 8 Simple Classification commands

#### 8.1 FALCON2 (no filter)

```
/usr/bin/time -v -- ./FALCON2 \
    meta -v -F -t 15 -l 47 -n 8 -x output_file.txt \
    reads.fq input-sequences.fna
```

#### 8.2 FALCON2 using the magnet functionality

```
/usr/bin/time -v -- ./FALCON2 \
    meta -mg -mf human_genome.fa -mt 0.50 \
    -v -F -t 15 -l 47 -n 8 -x output_file.txt \
    reads.fq \
    viruses.fa.
```

##### 8.3 Kraken2

```
/usr/bin/time -v -- kraken2 \
    -db kraken2_db/ --threads 8 \
    reads.fq > output_file.txt
```

##### 8.4 Centrifuge

```
/usr/bin/time -v -- centrifuge \
    -p 8 -x centrifuge_db -q reads.fq > output_file.txt
```

##### 8.5 CLARK-S

CLARK requires two steps: (1) classify reads, then (2) estimate abundances.

```
/usr/bin/time -v -- ./classify_metagenome.sh \
    -O reads.fq \
    -R classified.ready.csv \
    --light \
    -n 8
```

```
/usr/bin/time -v -- ./estimate_abundance.sh \
    -D CLARK_db/ \
    -F classified.ready.csv \
    > output_file.txt
```

#### 9 Benchmark Results

The full benchmark results, including items not cited in the main text. Paths are relative to the repository root. Classification performance varies with sequencing depth, read length, and deamination. AUROC, AUPRC, and $F_1$ are reported both after and before trimming, under contaminated and uncontaminated conditions.

Factor plots emphasizing precision–recall trade-offs under class imbalance, comparing trimming status and contamination conditions.

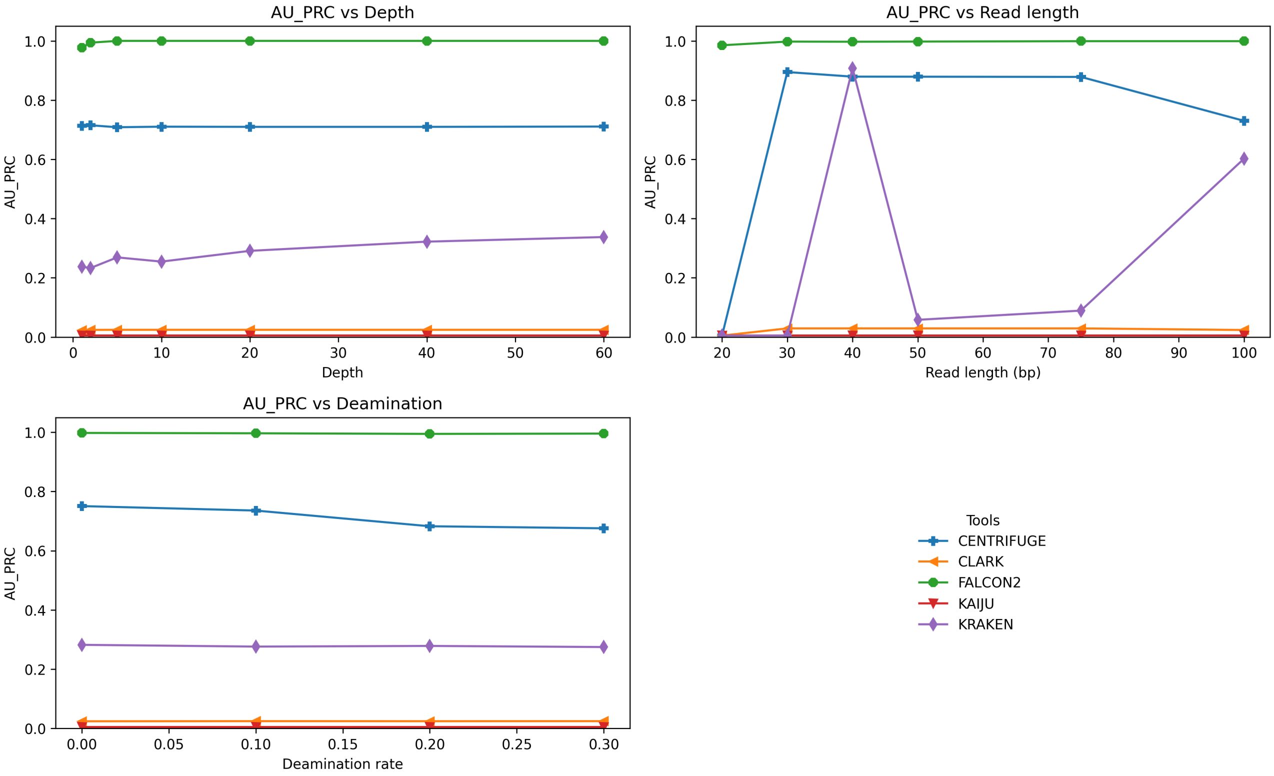


Figure S2: AUPRC after trimming, with contamination, across depth, read length, and deamination.

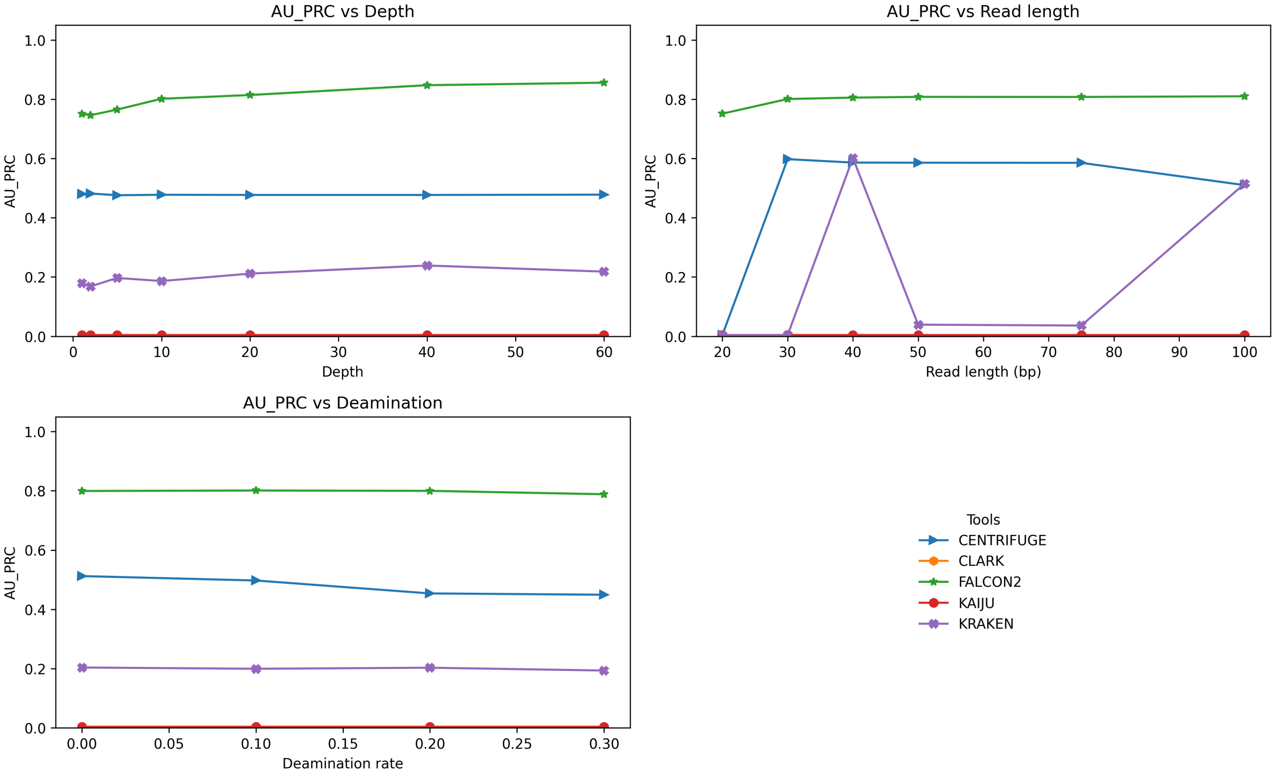


Figure S3: AUPRC after trimming, without contamination, across depth, read length, and deamination.

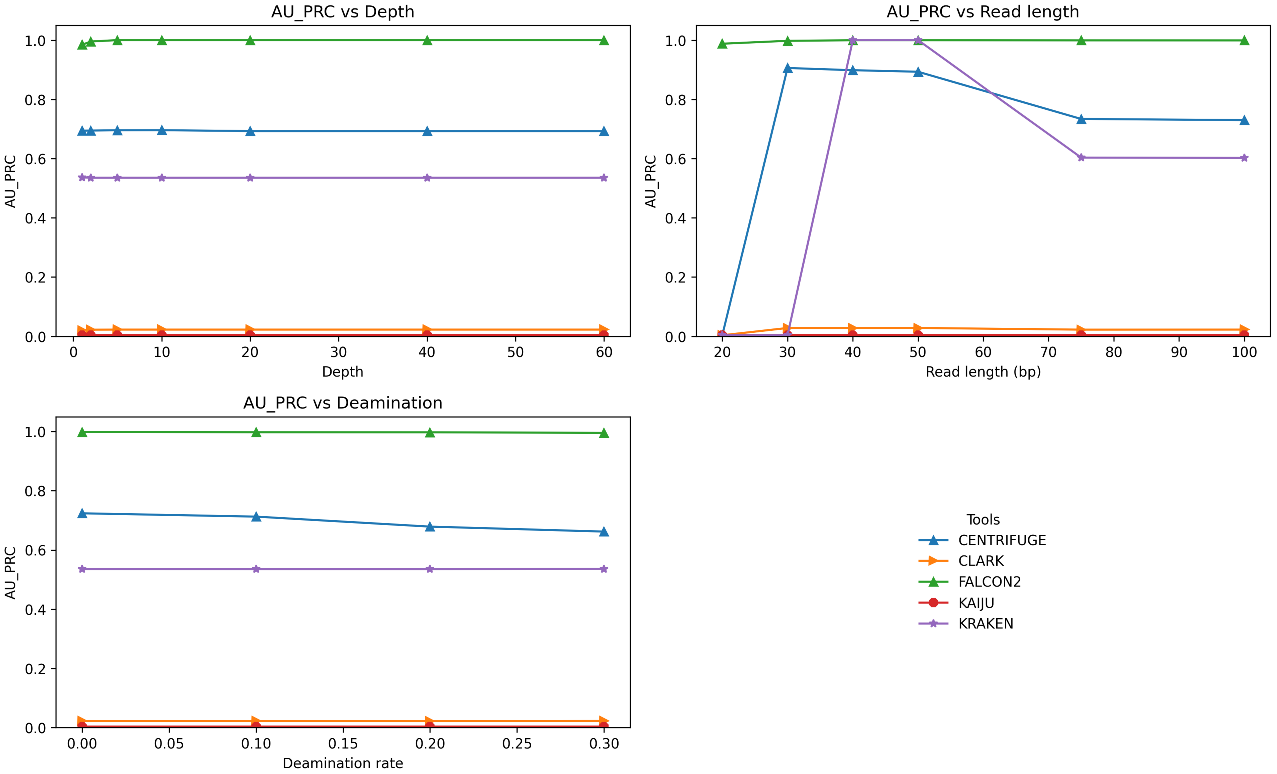


Figure S4: AUPRC before trimming, with contamination, across depth, read length, and deamination.

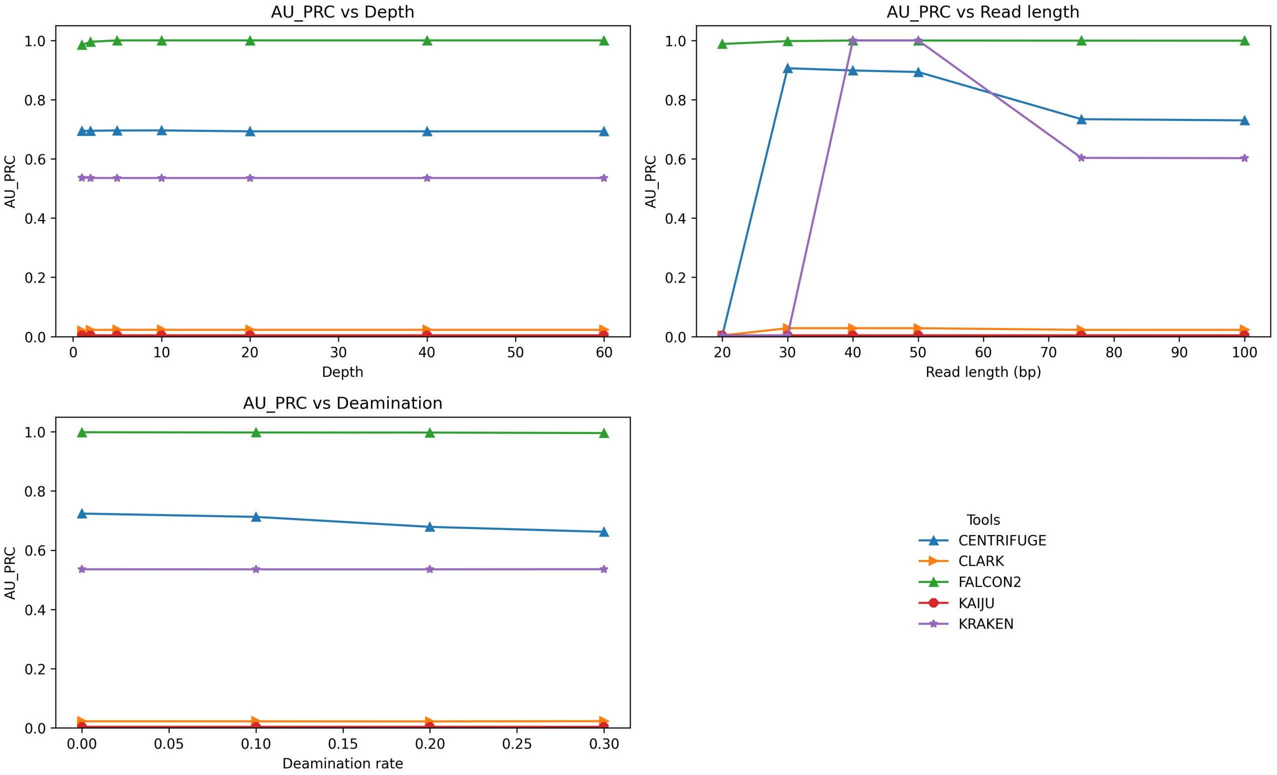


Figure S5: AUPRC before trimming, without contamination, across depth, read length, and deamination.

#### 9.1 F1-Score

Factor plots highlighting the balance between precision and recall across factors, with and without trimming and contamination.

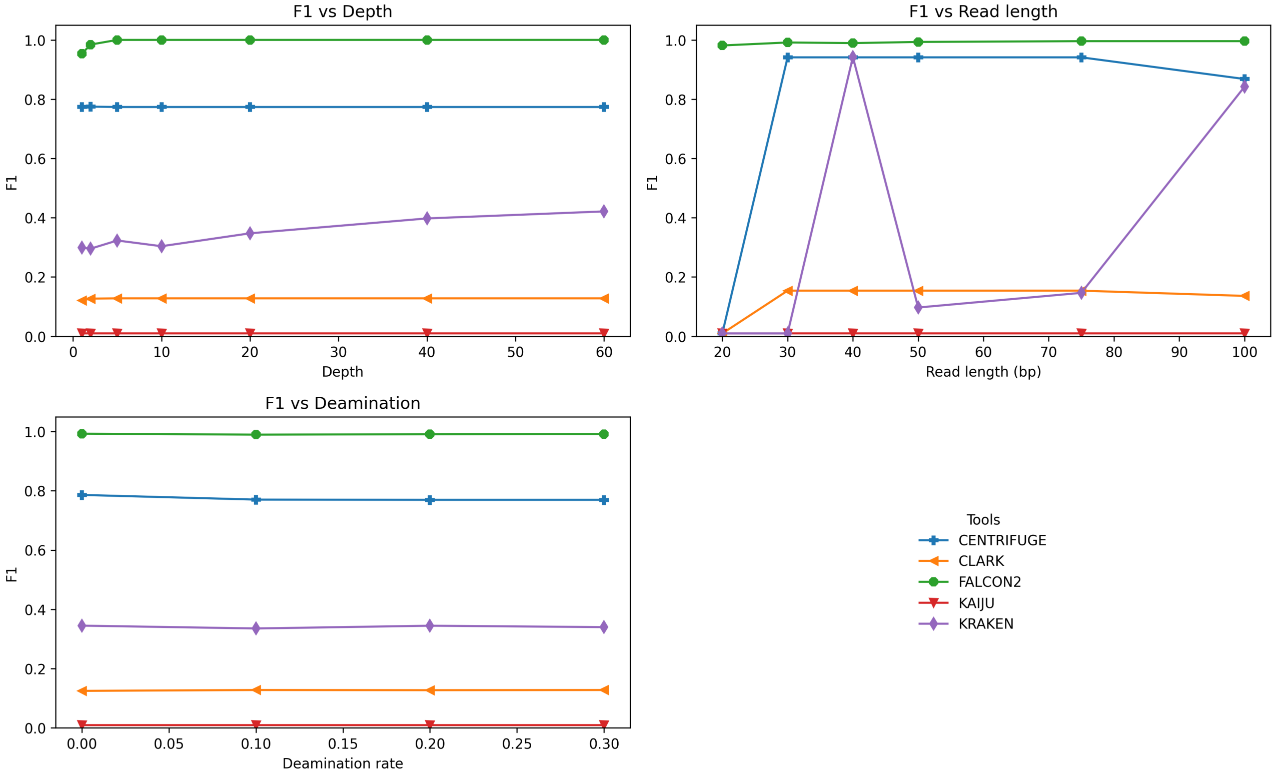


Figure S6: $F_1$-score after trimming, with contamination, across depth, read length, and deamination.

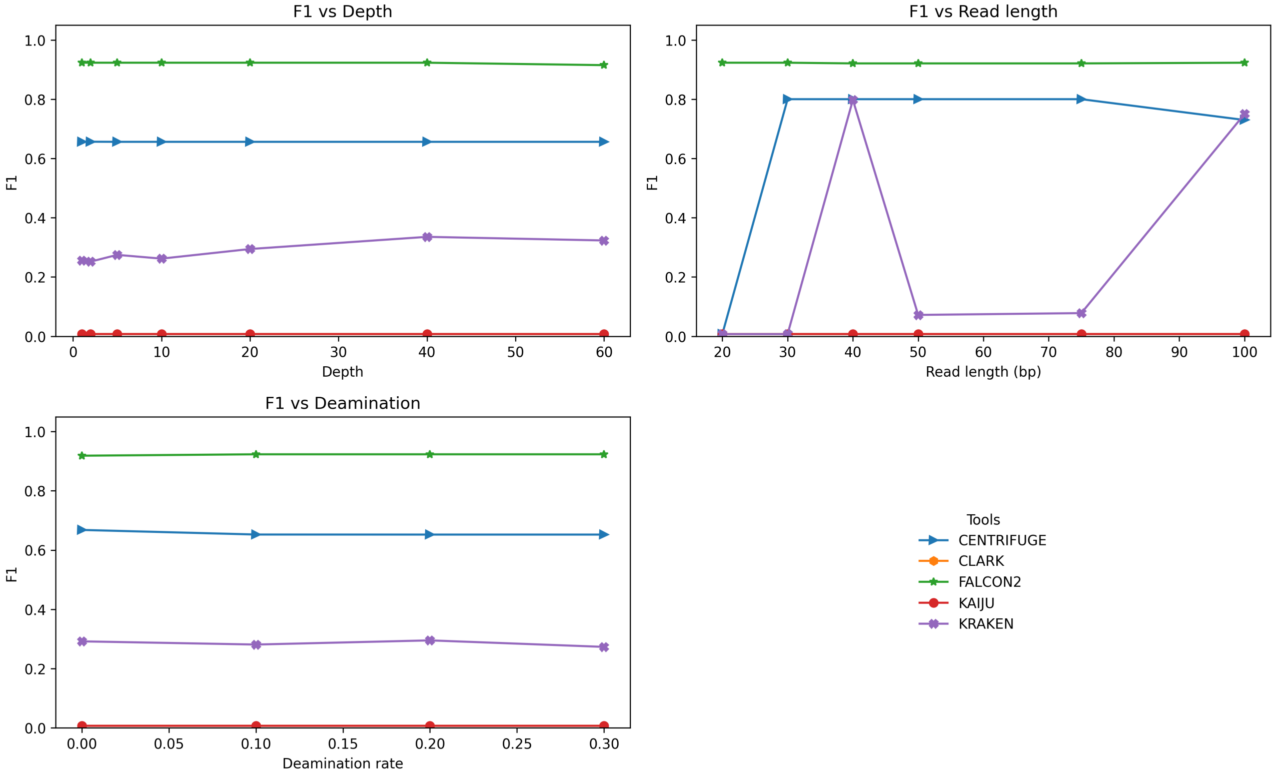


Figure S7: $F_1$-score after trimming, without contamination, across depth, read length, and deamination.

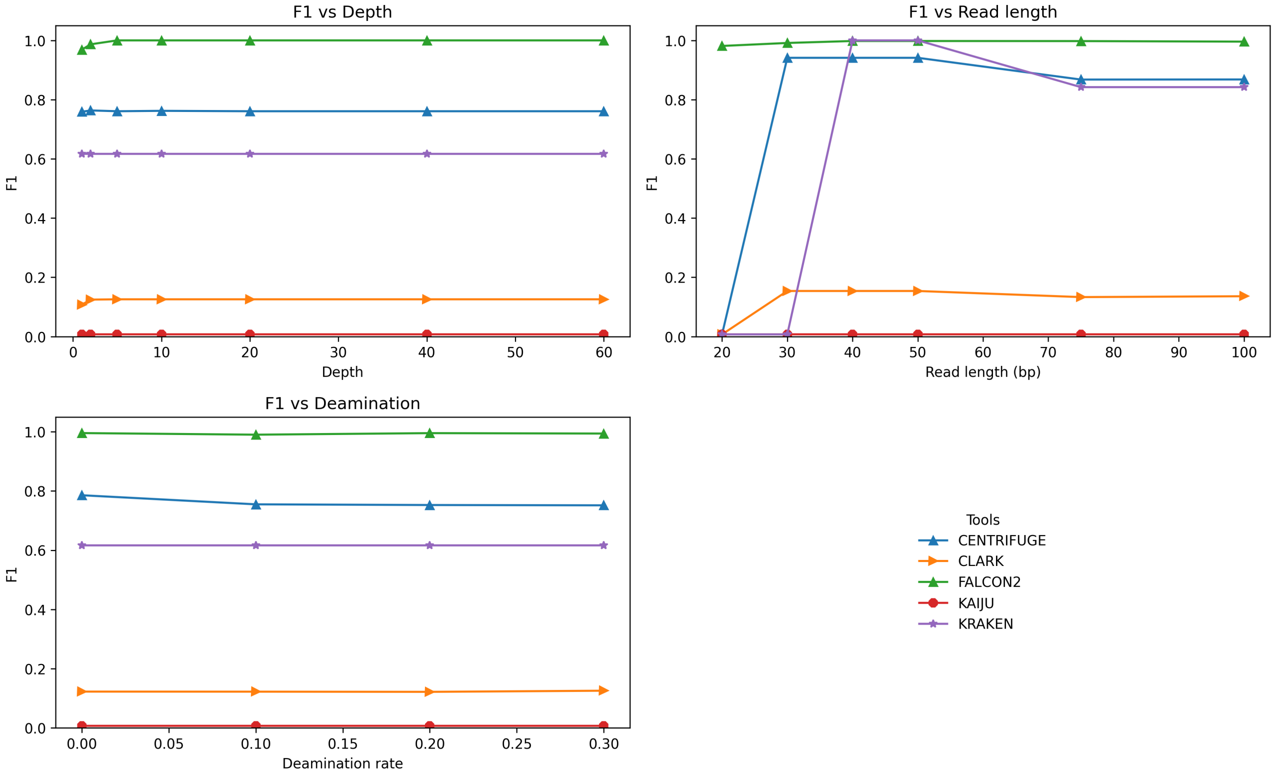


Figure S8: $F_1$-score before trimming, with contamination, across depth, read length, and deamination.

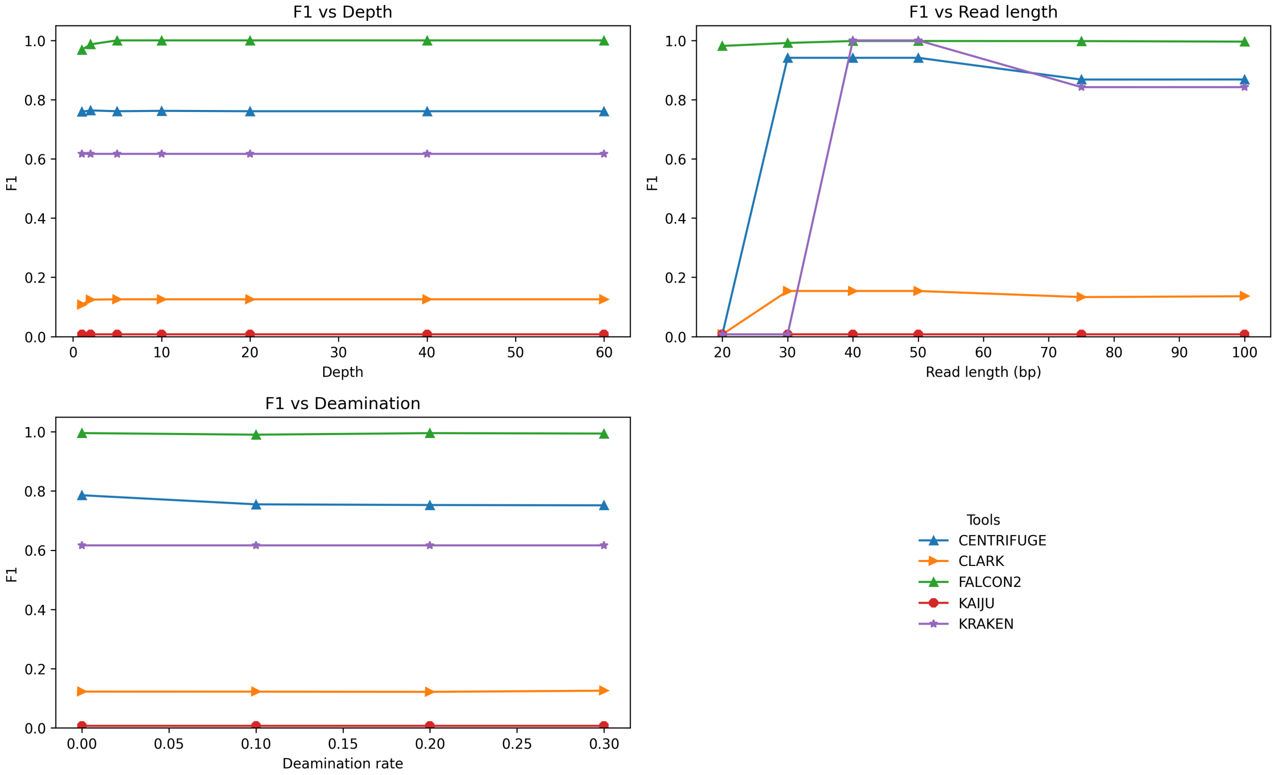


Figure S9: $F_1$-score before trimming, without contamination, across depth, read length, and deamination.

##### 9.1.1 AUROC

Factor plots showing global separability under different experimental factors, comparing trimmed vs. untrimmed and with vs. without contamination.

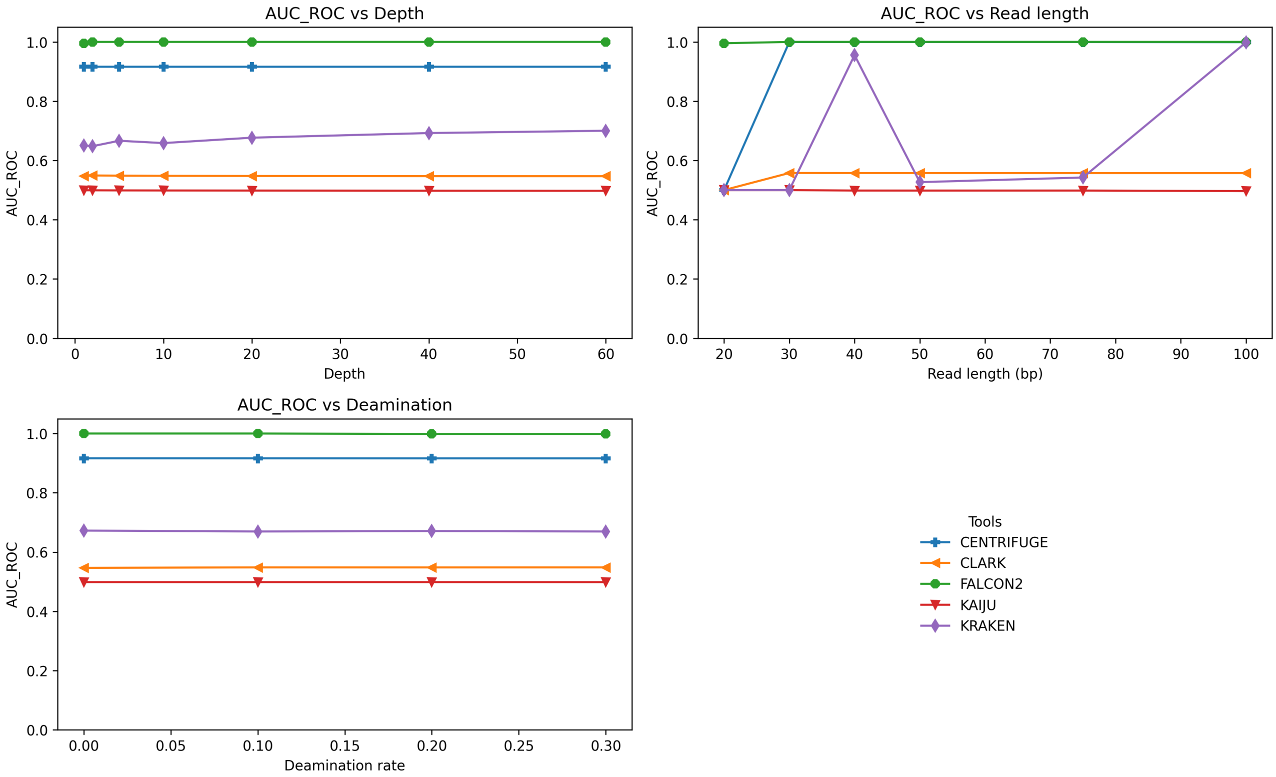


Figure S10: AUROC after trimming, with contamination, across depth, read length, and deamination.

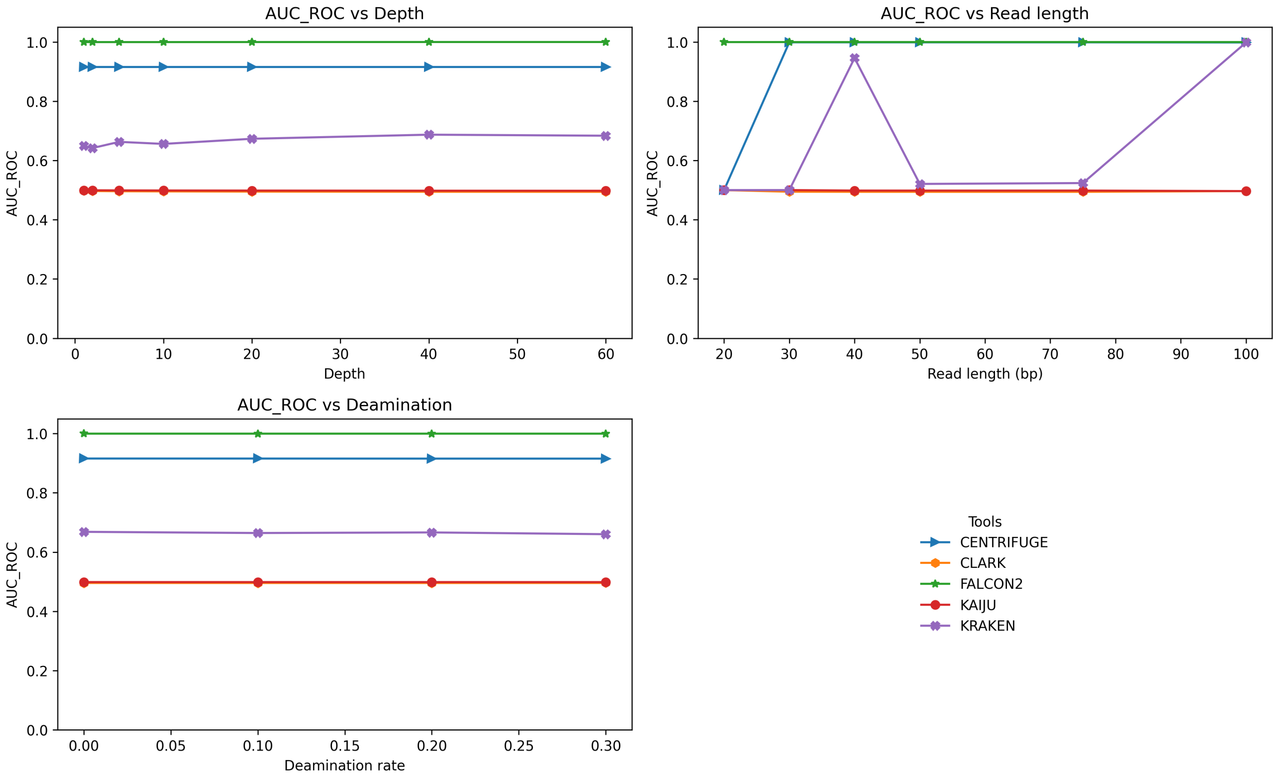


Figure S11: AUROC after trimming, without contamination, across depth, read length, and deamination.

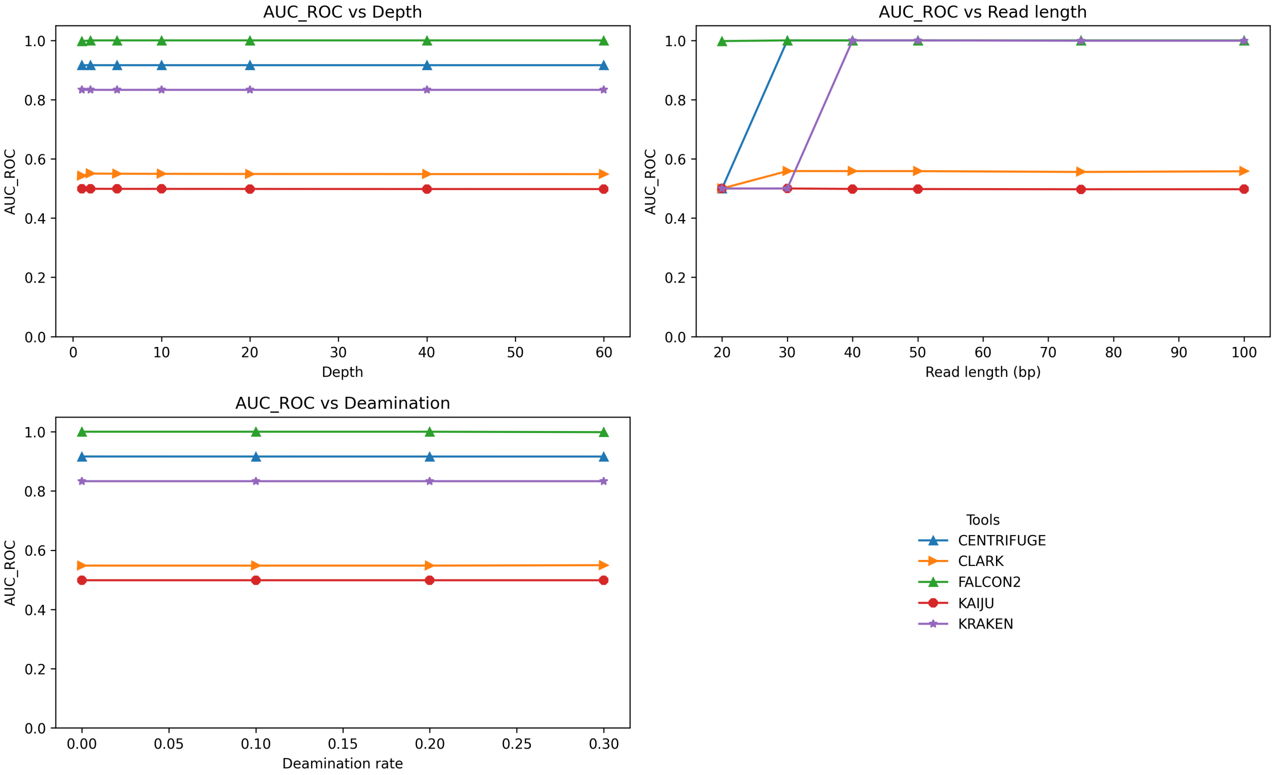


Figure S12: AUROC before trimming, with contamination, across depth, read length, and deamination.

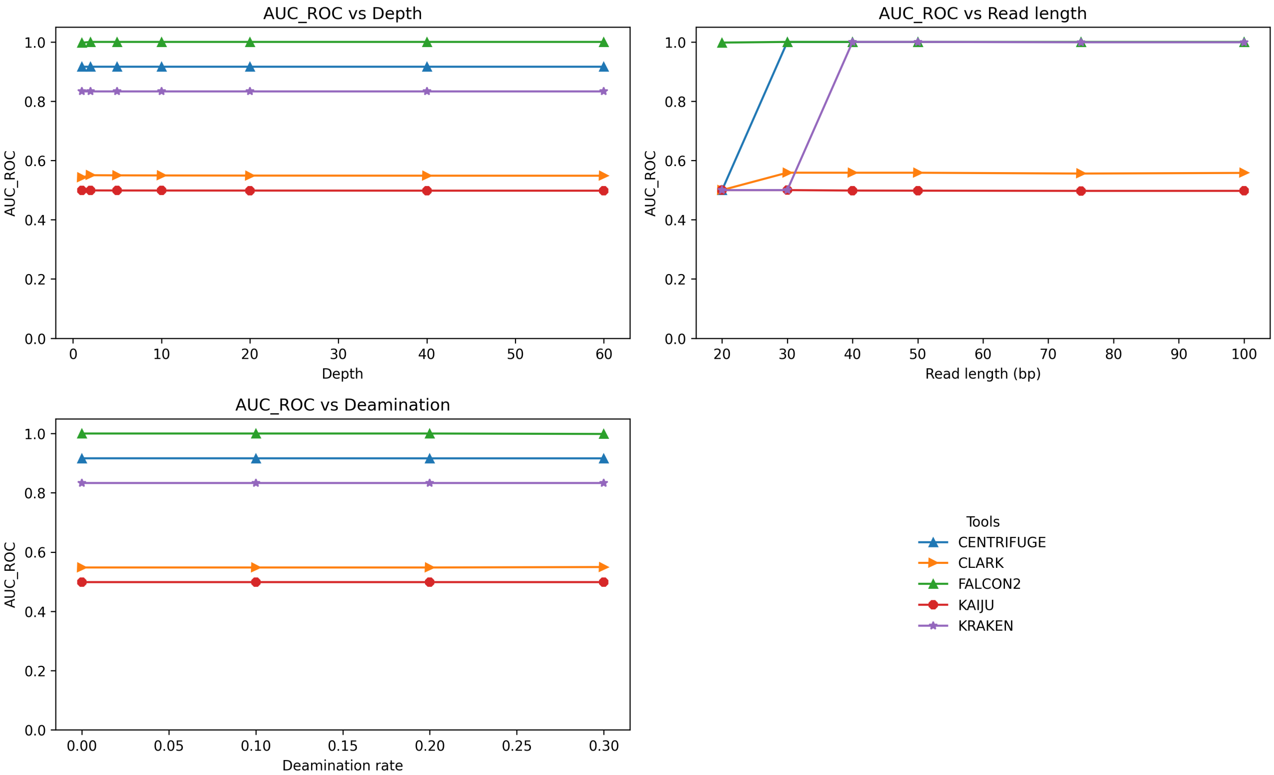


Figure S13: AUROC before trimming, without contamination, across depth, read length, and deamination.

### 10 mapDamage Profiles

Diagnostic damage patterns used to evaluate trimming effects across deamination levels.

#### 10.1 After Trimming

Profiles at $\delta \in \{0, 0.1, 0.2, 0.3\}$ showing residual damage signatures post-trimming.

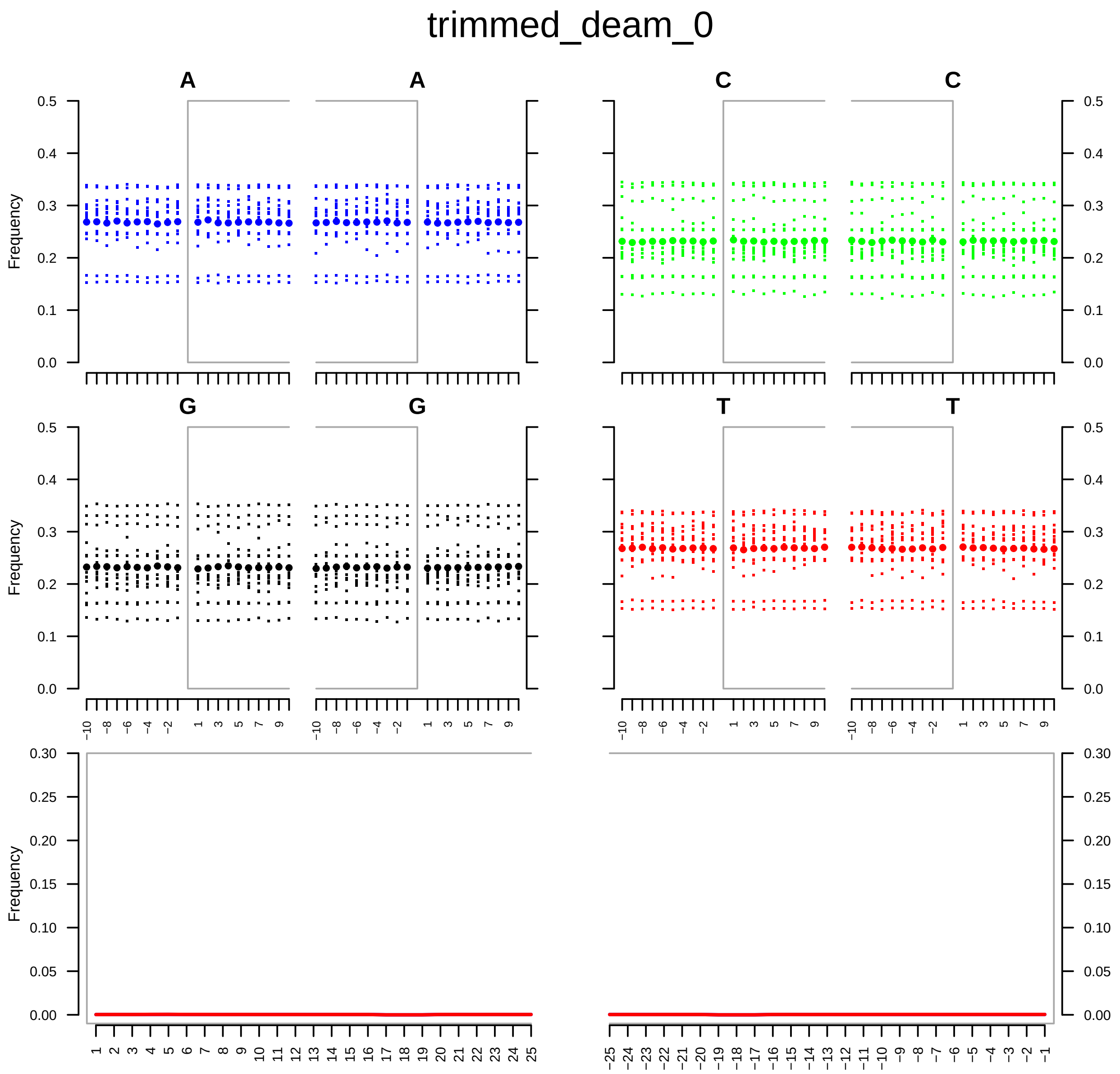


Figure S14: mapDamage profiles after trimming at deamination $\delta = 0.0$.

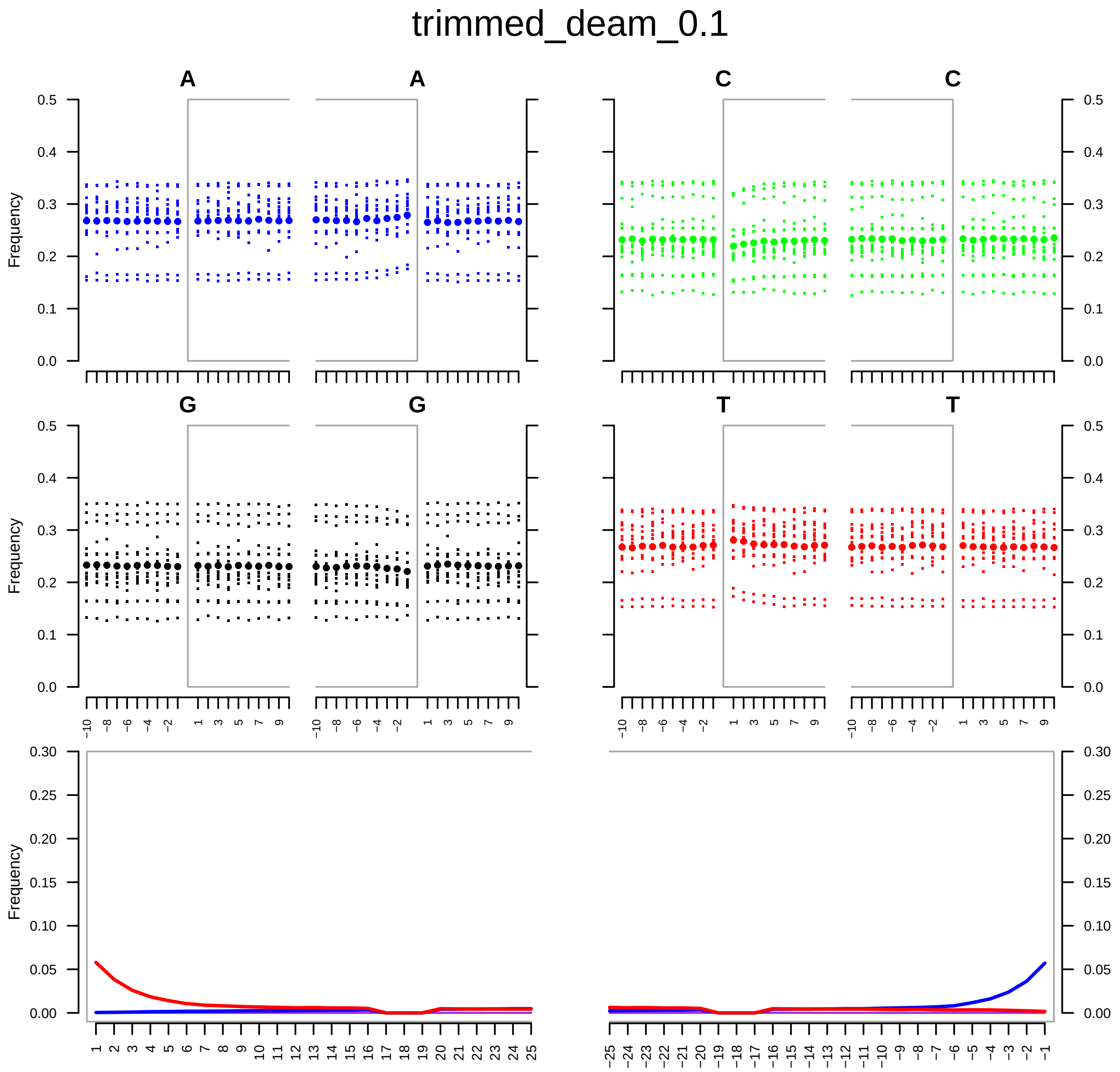


Figure S15: mapDamage profiles after trimming at deamination $\delta = 0.1$.

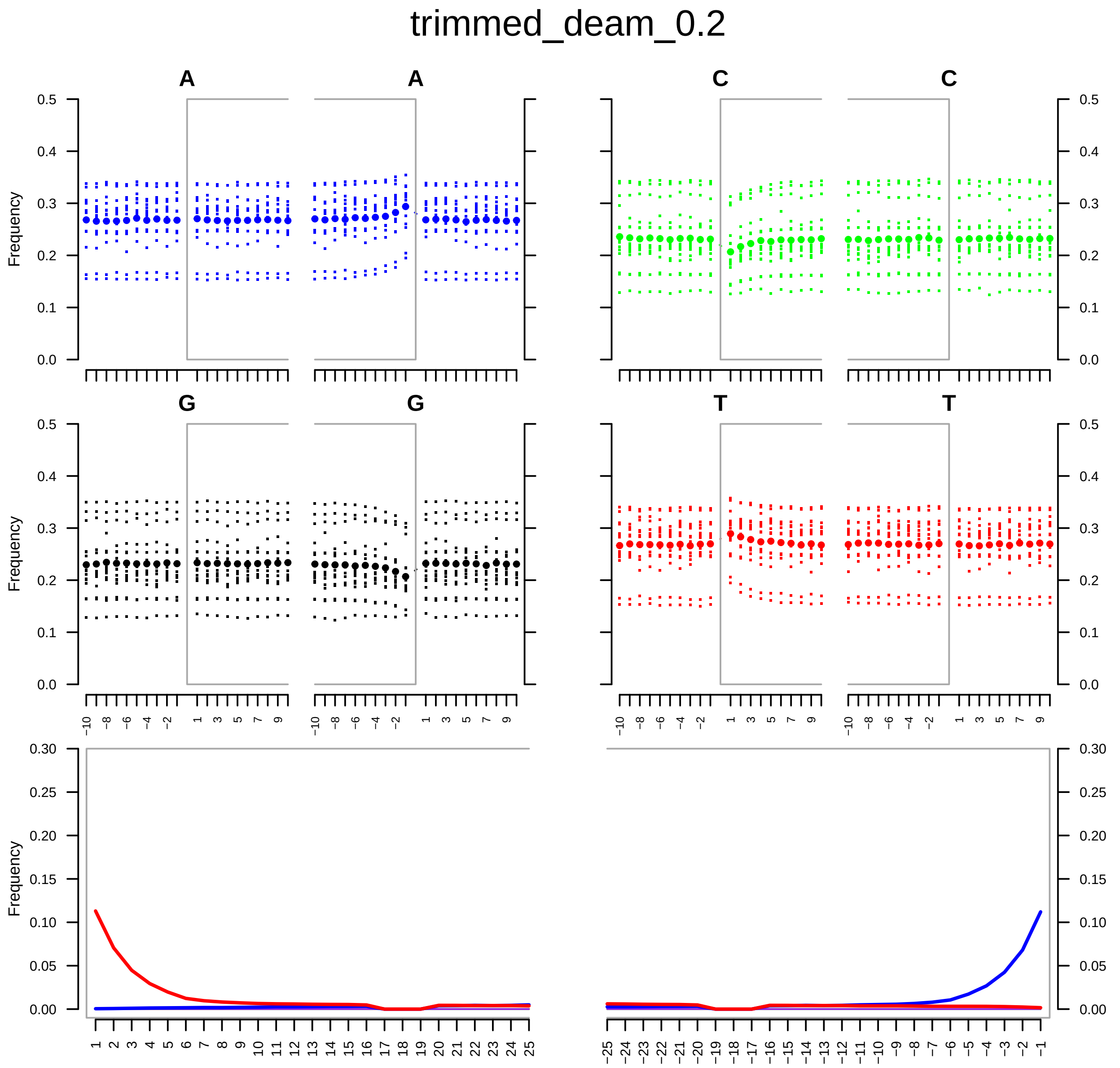


Figure S16: mapDamage profiles after trimming at deamination $\delta = 0.2$.

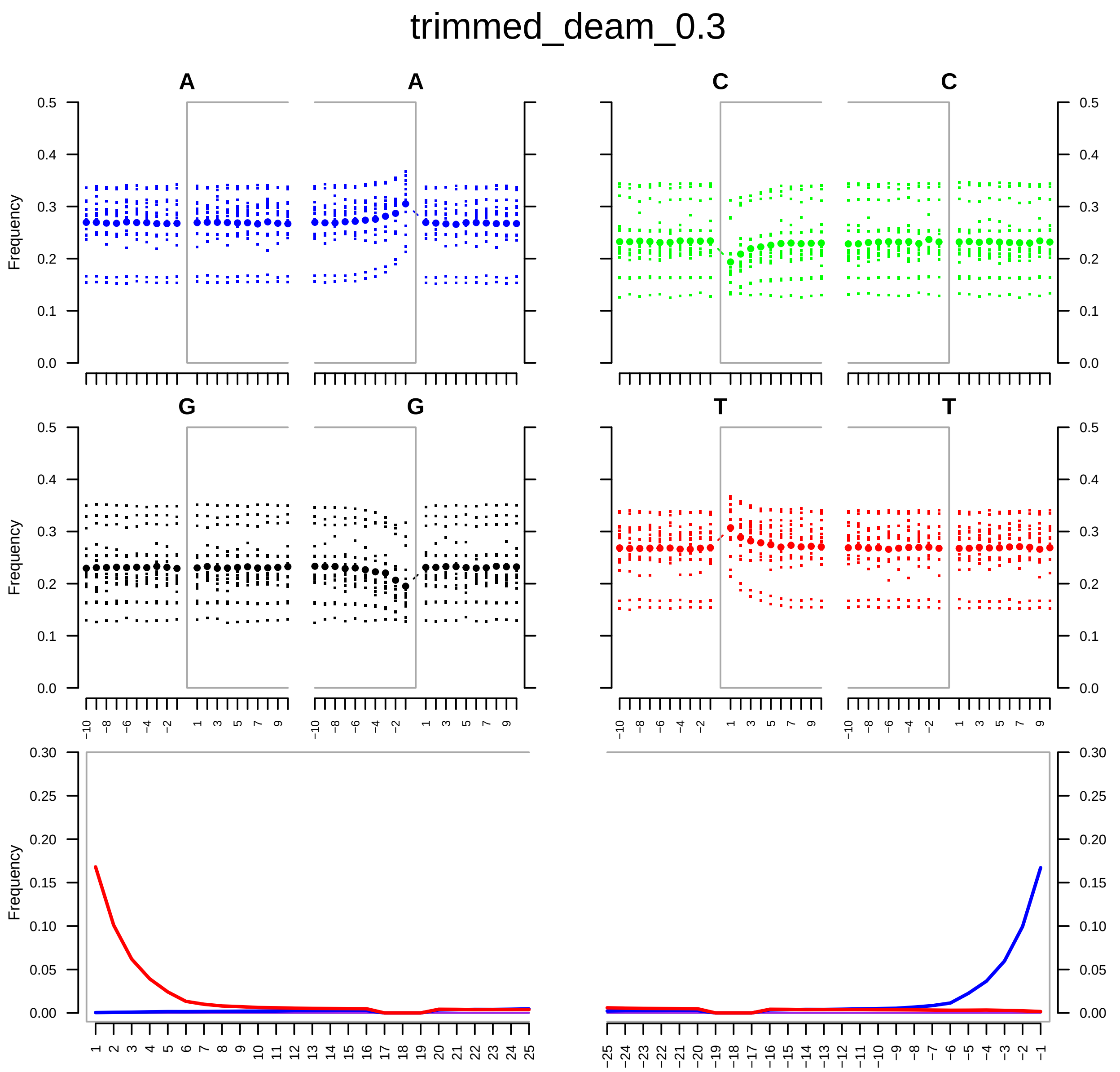


Figure S17: mapDamage profiles after trimming at deamination $\delta = 0.03$.

##### 10.1.1 Before Trimming

Profiles at $\delta \in \{0, 0.1, 0.2, 0.3\}$ showing raw damage patterns prior to trimming.

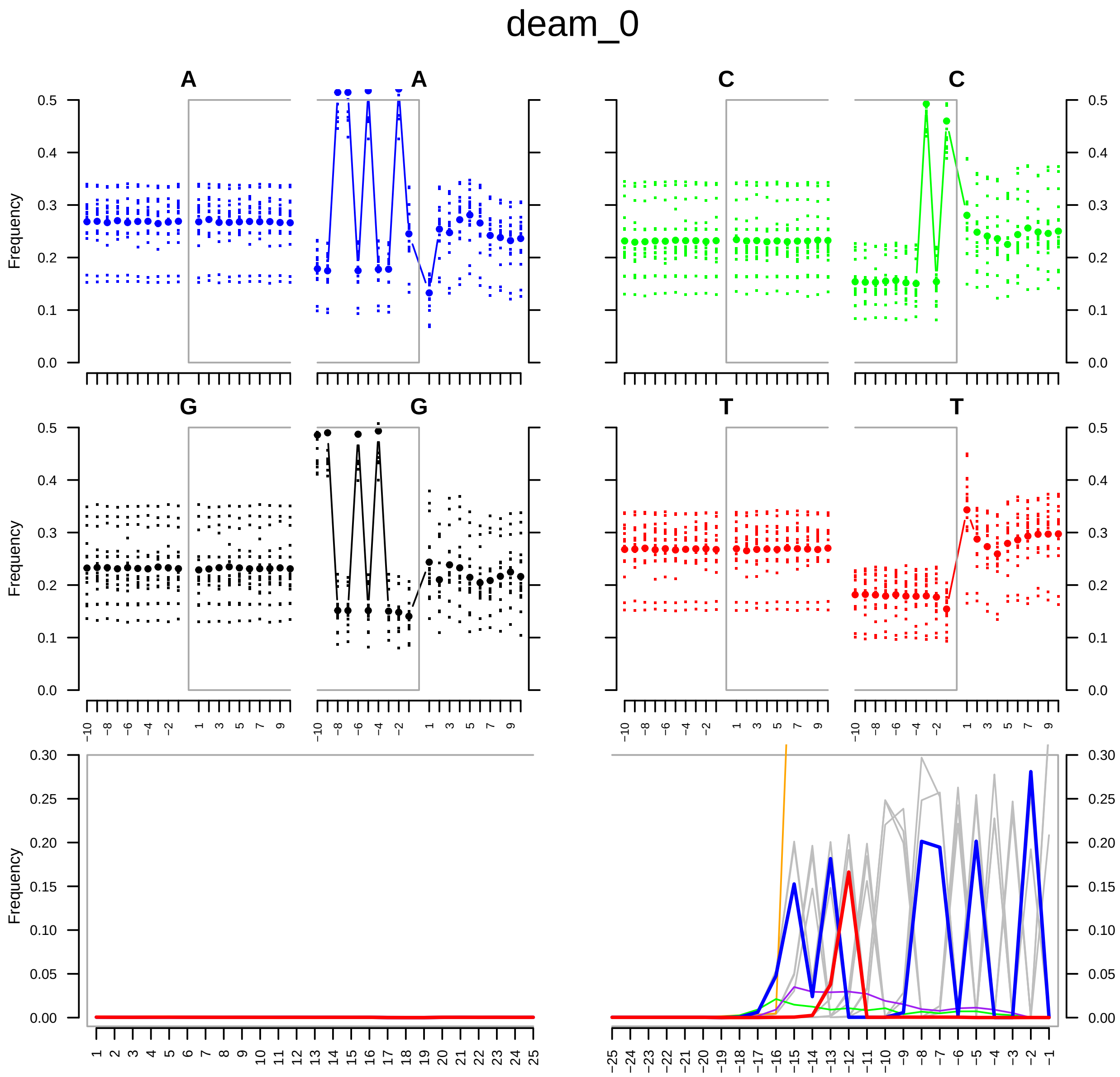


Figure S18: mapDamage profiles for untrimmed reads $\delta = 0.0$.

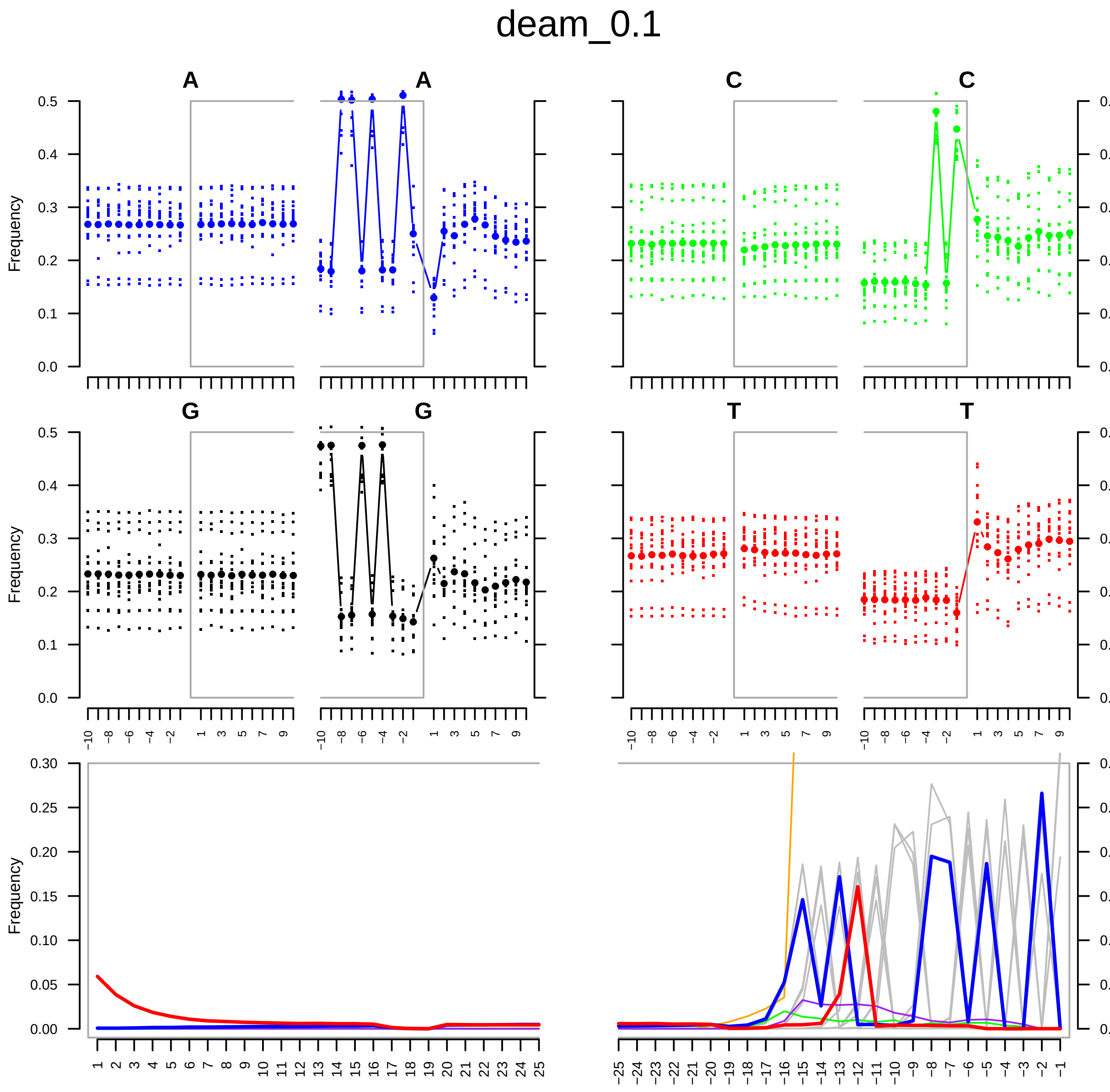


Figure S19: mapDamage profiles for untrimmed reads $\delta = 0.1$.

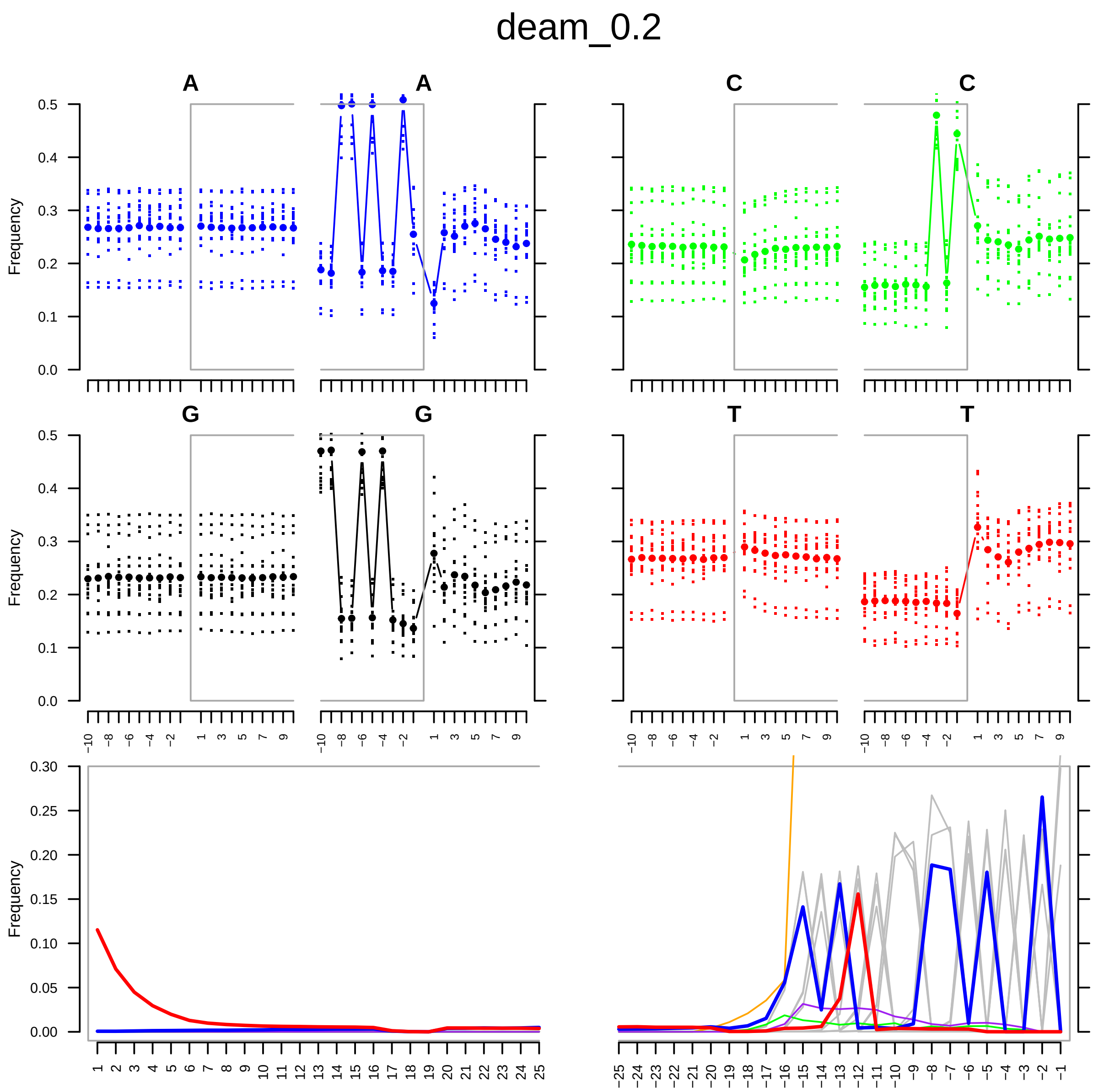


Figure S20: mapDamage profiles for untrimmed reads $\delta = 0.2$.

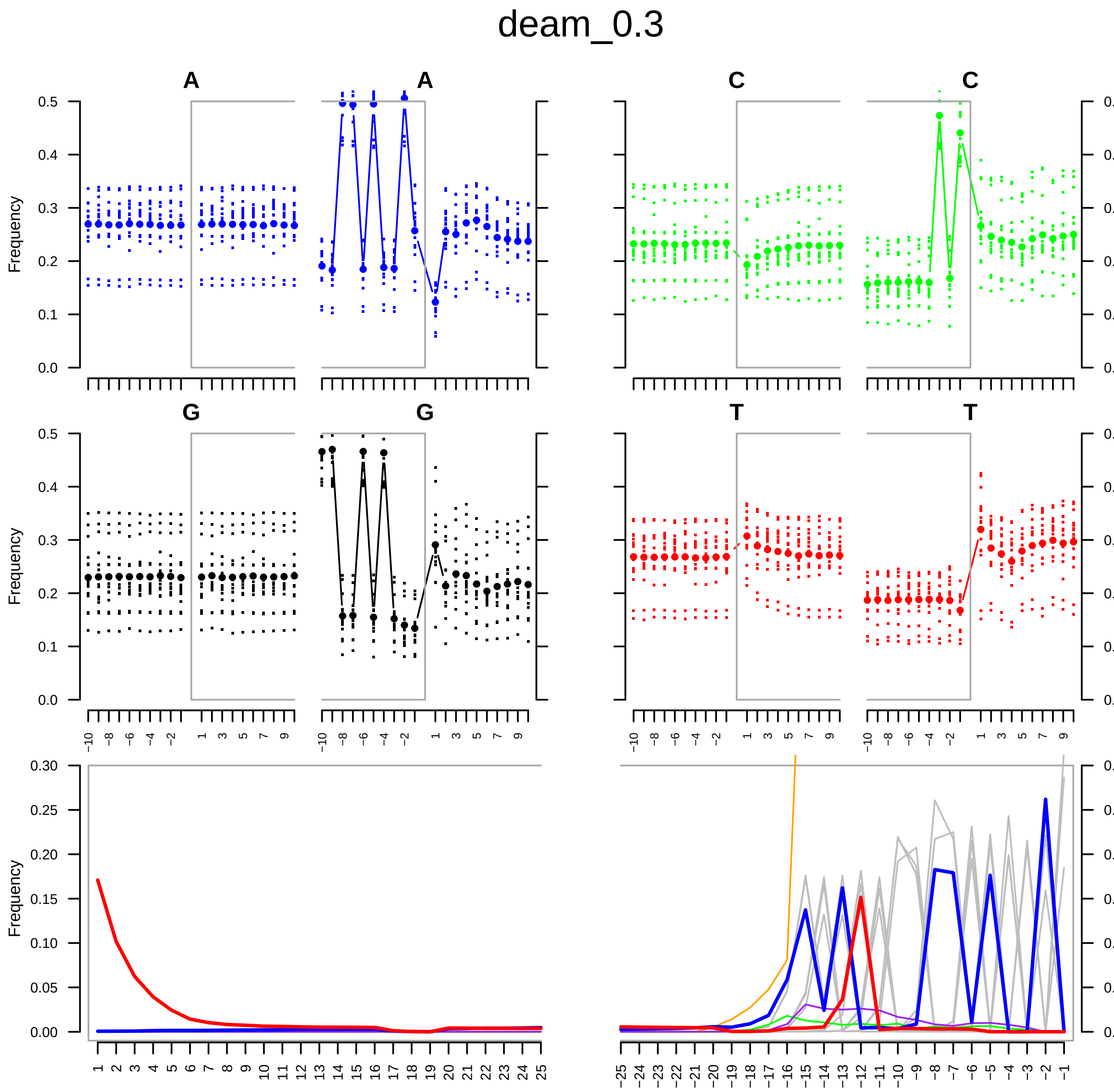


Figure S21: mapDamage profiles for untrimmed reads $\delta = 0.3$.

### 11 Metric Slices (Trimmed, With Contamination)

'Slice' views where one factor varies while the others are fixed, isolating its effect under contamination.

#### 11.1 AUPRC Slices

AUPRC under the same three slice settings as AUROC to examine precision–recall sensitivity under contamination.

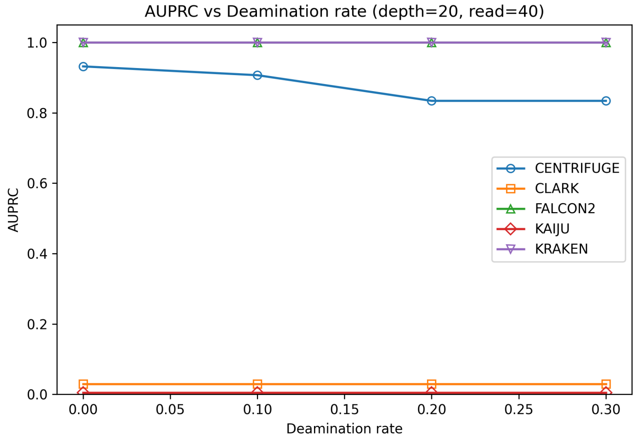


Figure S22: AUPRC after trimming, with contamination; varying deamination ($\delta$) at fixed depth = 20× and read length = 40 bp.

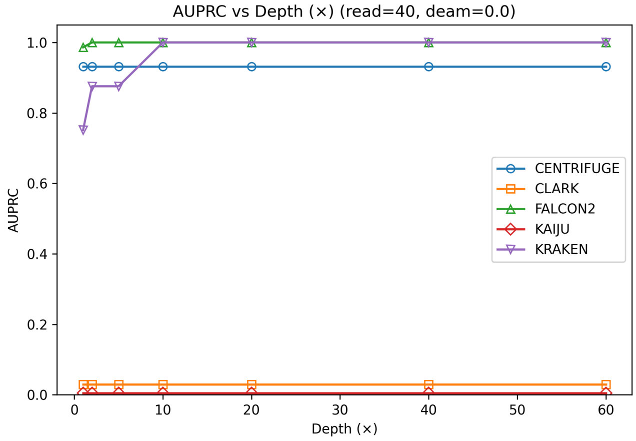


Figure S23: AUPRC after trimming, with contamination; varying depth at fixed read length = 40 bp and deamination $\delta = 0.0$.

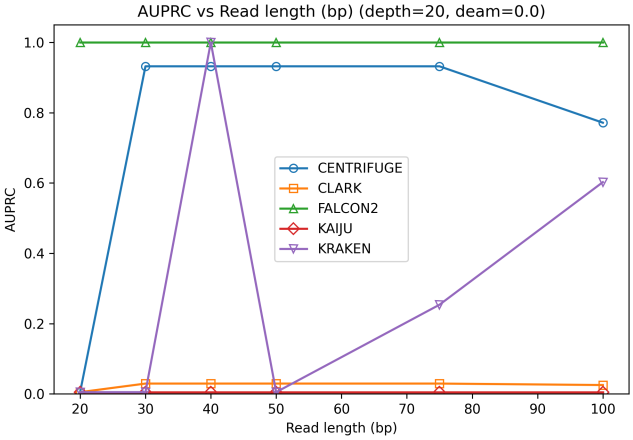


Figure S24: AUPRC after trimming, with contamination; varying read length at fixed depth = 20× and deamination $\delta = 0.0$.

###### 11.1.1 F1 Slices

$F_1$ under the same slice settings, highlighting the precision/recall balance when only one factor changes.

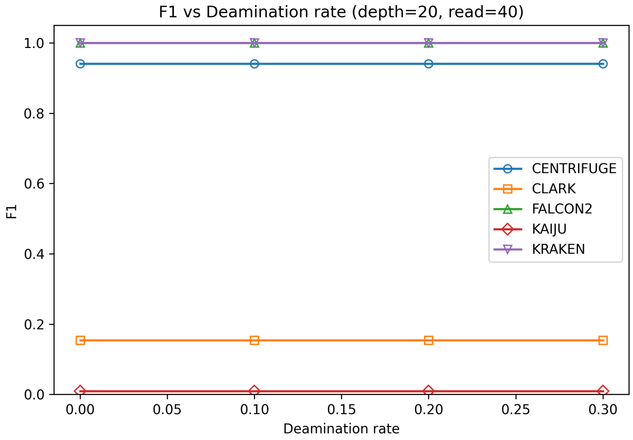


Figure S25: $F_1$-score after trimming, with contamination; varying deamination ($\delta$) at fixed depth = 20× and read length = 40 bp.

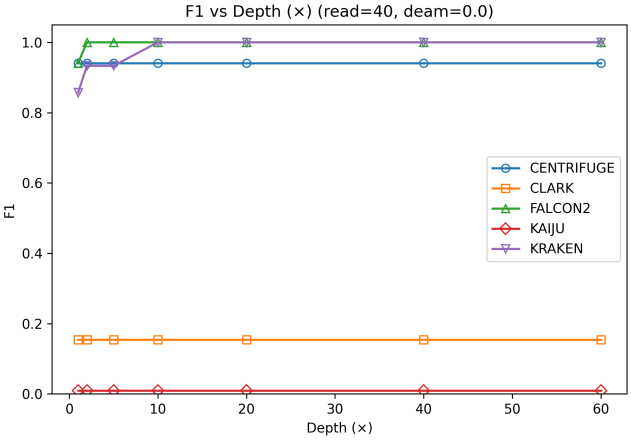


Figure S26: $F_1$-score after trimming, with contamination; varying depth at fixed read length = 40 bp and deamination $\delta = 0.0$.

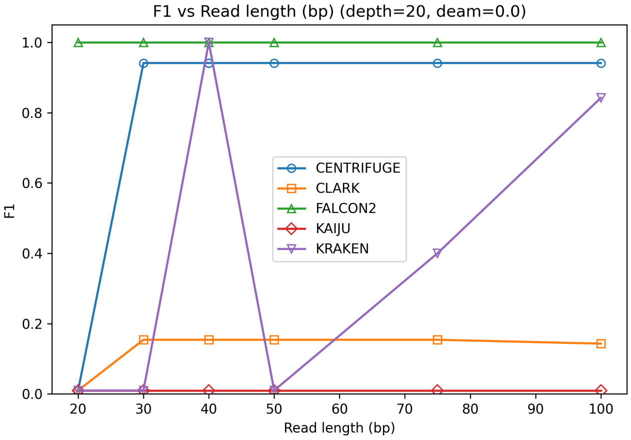


Figure S27: $F_1$-score after trimming, with contamination; varying read length at fixed depth = 20× and deamination $\delta = 0.0$.

##### 11.1.2 AUROC Slices

AUROC as a function of (i) deamination at fixed depth 20× and read length 40 bp, (ii) depth at fixed read length 40 bp and $\delta = 0$, and (iii) read length at fixed depth 20× and $\delta = 0$.

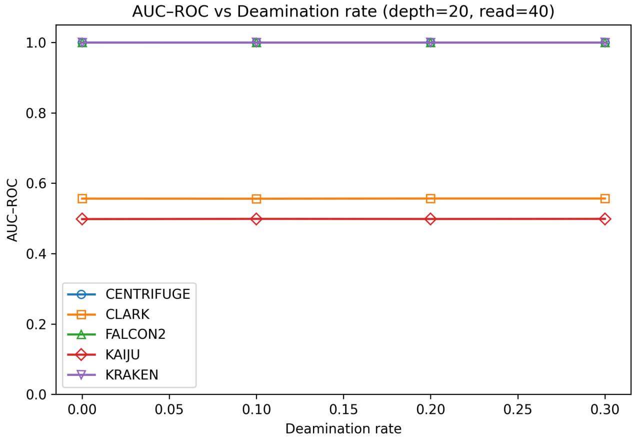


Figure S28: AUROC after trimming, with contamination; varying deamination ($\delta$) at fixed depth = 20× and read length = 40 bp.

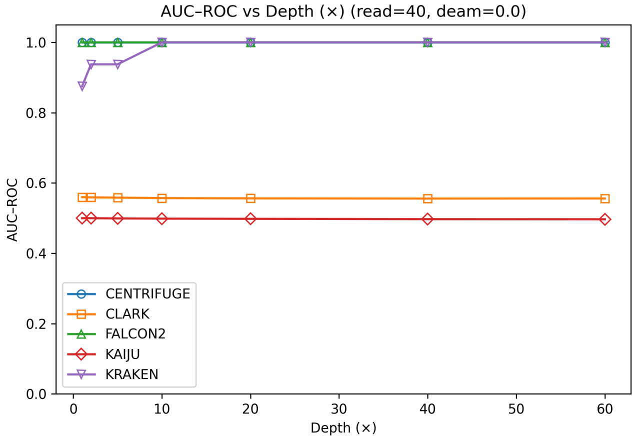


Figure S29: AUROC after trimming, with contamination; varying depth at fixed read length = 40 bp and deamination $\delta = 0.0$.

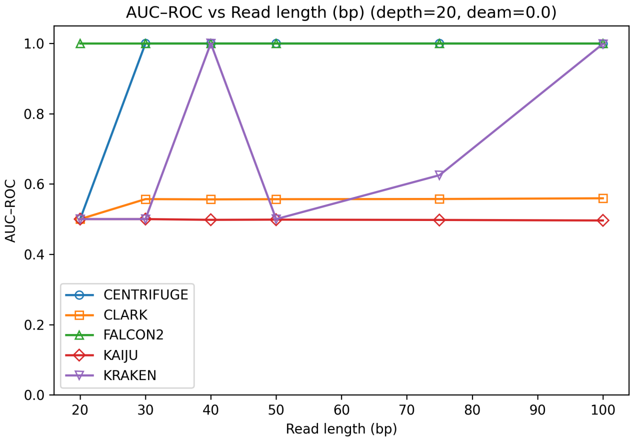


Figure S30: AUROC after trimming, with contamination; varying read length at fixed depth $= 20\times$ and deamination $\delta = 0.0$.

### 12 PR/ROC Curves (Trimmed, Without Contamination)

Method-by-method discrimination curves after trimming in clean conditions.

#### 12.1 Precision–Recall

PR curves for Centrifuge, FALCON2, and Kraken2 to assess performance under class imbalance.

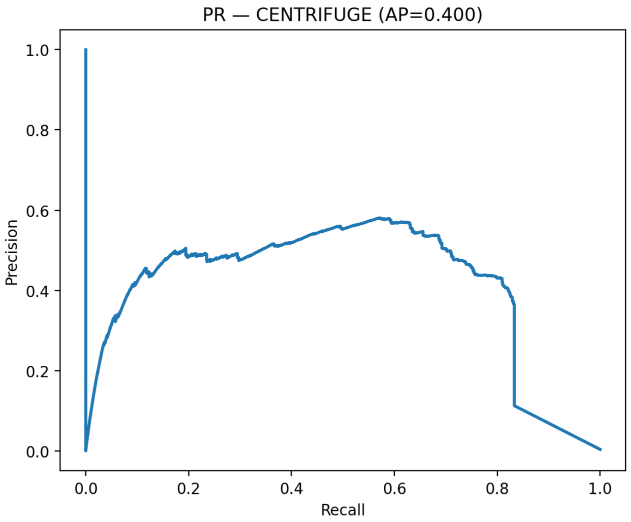


Figure S31: Precision–Recall curves after trimming, without contamination (Centrifuge).

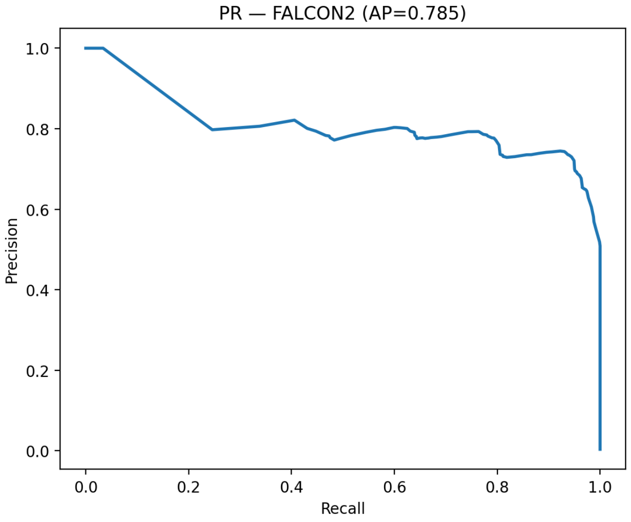


Figure S32: Precision–Recall curves after trimming, without contamination (FALCON2).

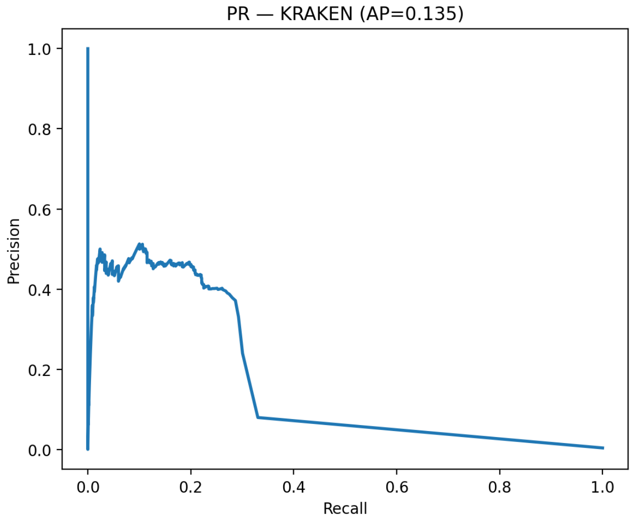


Figure S33: Precision–Recall curves after trimming, without contamination (Kraken2).

###### 12.1.1 ROC

ROC curves for the same methods, providing a threshold-independent view of separability without contamination.

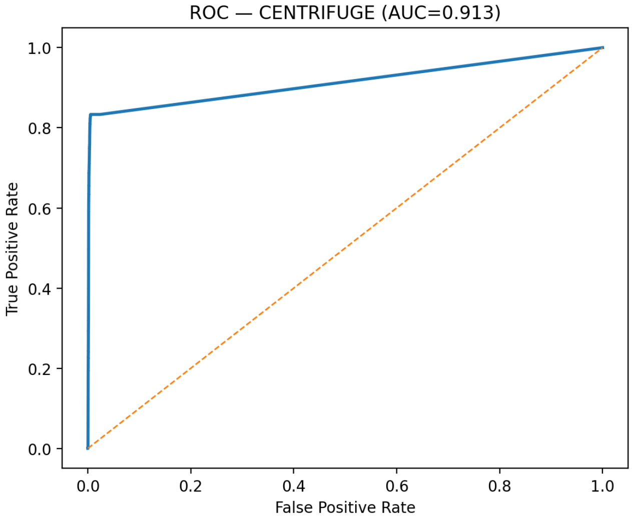


Figure S34: ROC curves after trimming, without contamination (Centrifuge).

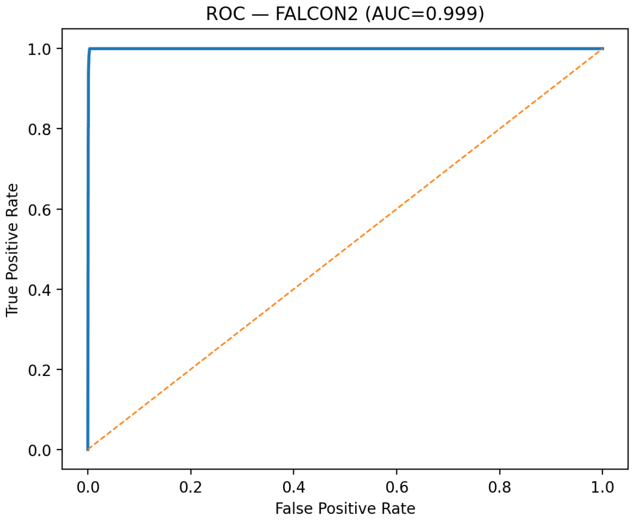


Figure S35: ROC curves after trimming, without contamination (FALCON2).

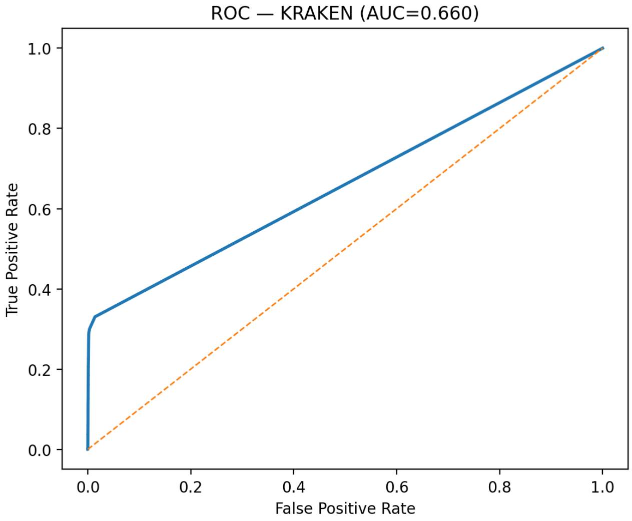


Figure S36: ROC curves after trimming, without contamination (Kraken2).

### 13 PR/ROC Curves (Trimmed, With Contamination)

Method-by-method discrimination curves after trimming in contaminated conditions.

#### 13.1 Precision–Recall

PR curves for Centrifuge, FALCON2, and Kraken2, highlighting robustness to contamination.

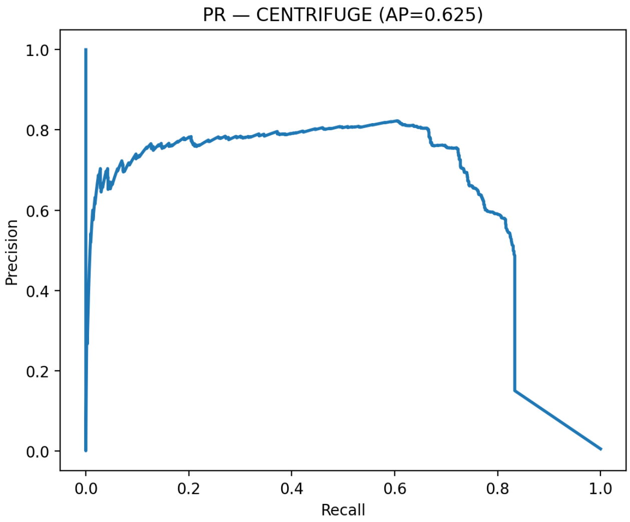


Figure S37: Precision–Recall curves after trimming, with contamination (Centrifuge).

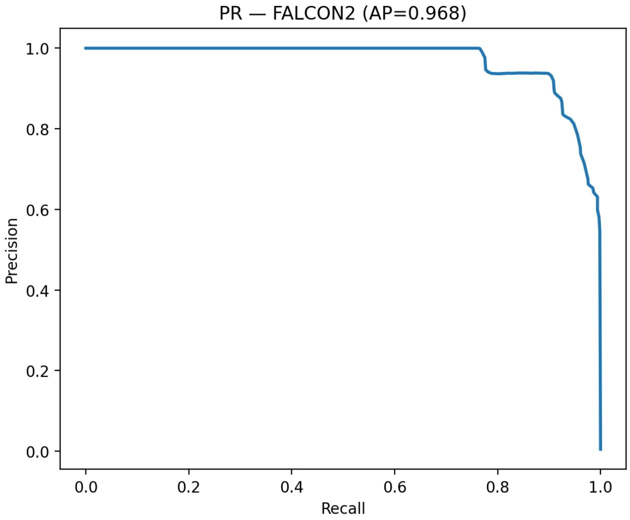


Figure S38: Precision–Recall curves after trimming, with contamination (FALCON2).

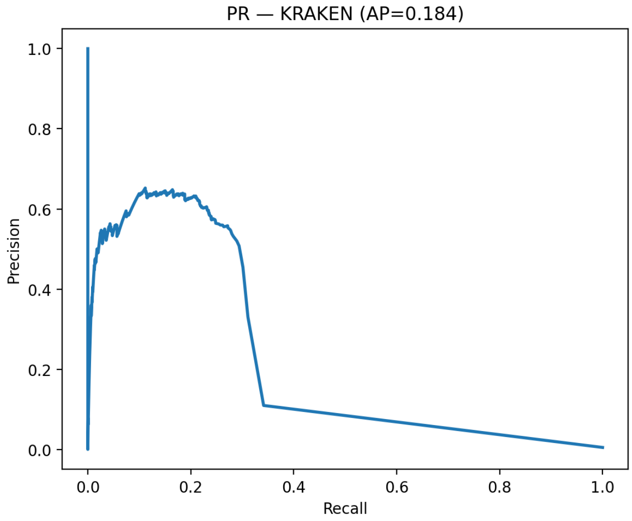


Figure S39: Precision–Recall curves after trimming, with contamination (Kraken2).

###### 13.1.1 ROC

ROC curves for the same methods, showing how contamination impacts true/false positive trade-offs.

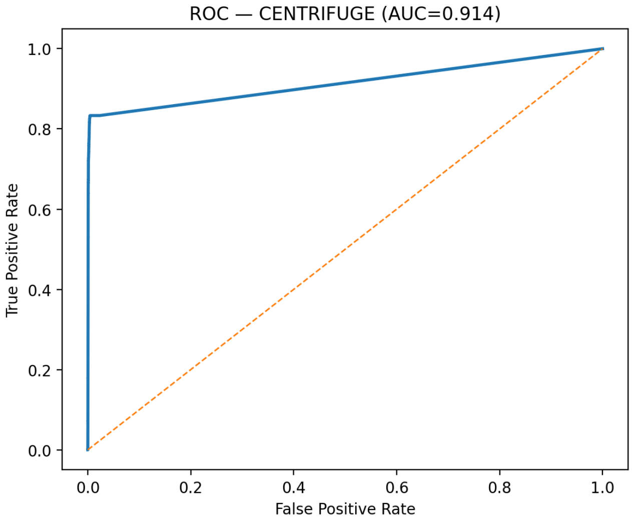


Figure S40: ROC curves after trimming, with contamination (Centrifuge).

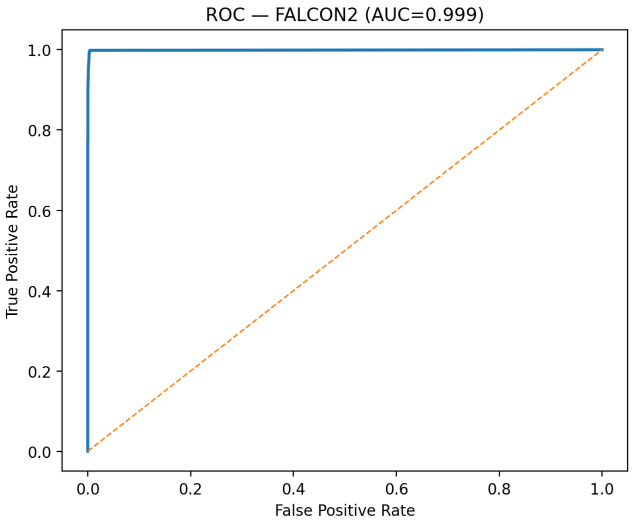


Figure S41: ROC curves after trimming, with contamination (FALCON2).

### 14 Archive

We archive the environment file (`env.yml`), timing and memory outputs from `/usr/bin/time -v`, per-condition metric CSVs, and the figure-generation scripts, and make them available under the persistent DOI `10.5281/zenodo.17291214`

### 15 The FALCON2 tool

#### Overview

**FALCON2 is an ultra-fast method to infer metagenomic composition of sequenced reads.** It measures **similarity between any FASTQ/FASTA file** against **any multi-FASTA database** (e.g., all complete NCBI genomes). It supports single and paired-end reads (Illumina MiSeq/HiSeq/Novaseq, IonTorrent, etc.), uses **relative data compression**, and employs **variable multi-threading** without multiplying memory per thread. Beyond global composition, FALCON2 can **locate local similarity regions** and provides subcommands to *filter* (`filter`), *visualize* (`fvisual`), *analyze inter-similarity* (`inter`), and *visualize inter-similarities* (`ivisual`).

#### 15.1 Installation

##### 15.1.1 Manual (from source)

```
git clone https://github.com/cobilab/falcon.git
cd falcon/src/
cmake .
make
cp FALCON2 ../../
cd ../../
```

*Requirement:* CMake.

### 16 Demo

Find a quick demo video in the repository (see README). To reproduce a simple test (search top 15 similar viruses in bundled reads):

```
cd test
gunzip reads.fq.gz
gunzip VDB.fa.gz
./FALCON2 meta -v -F -t 15 -l 47 -x top.txt reads.fq VDB.fa
```

The output `top.txt` will list the most similar references (e.g., Zaire Ebolavirus in this example).

### 17 Building a reference database

#### 17.1 Latest NCBI viral database (example)

```
# Download viral genomes from NCBI
./download_references_ncbi.sh viruses
```

#### 17.2 Download a ready-to-use viral DB

A prebuilt viral database is available online (see README). Download and use `VDB.fa-gz` with FALCON2.

### 18 Usage

#### 18.1 Main menu

To list available commands:

```
./FALCON2
# or
./FALCON2 -h
```

Output:

```
COMMANDS
  meta     - Infer metagenomic sample composition (main)
  filter   - Filter/segment regions identified by FALCON
  fvisual  - Visualize global/local similarities
  inter    - Evaluate inter-similarity between genomes
  ivisual  - Heatmap visualization of genome similarities

Use 'FALCON2 <command> -h' for help with a specific command.
```

#### 18.2 Metagenomic composition analysis (`meta`)

##### 18.2.1 Help

```
./FALCON2 meta -h
```

##### 18.2.2 Options (abridged)

```
Non-mandatory:
  -h, --help                 show help
  -F, --force                overwrite output files
  -V, --version              show version and exit
  -v, --verbose              verbose mode
  -Z, --local                compute database local similarity
  -s, --show                 show compression levels

  -l, --level <1..47>        compression level
  -p, --sample <rate>        subsampling (default: 1)
  -t, --top <num>            top-N to report (default: 20)
  -n, --nThreads <num>       threads (default: 2)

  -x, --output <file>        similarity top filename
  -y, --profile <file>       profile filename (-Z must be on)

  -S, --save-model           save models after learning
  -L, --load-model           load previously saved model
  -M, --model-file <file>    model filename
  -I, --model-info           show model info
  -T, --train-model          train model only (no inference)

Mandatory:
  [FASTQ[:FASTQ...]]         metagenomic reads (FASTQ)
  [FASTA[:FASTA...]]         database (multi-FASTA)

MAGNET integration:
  -mg, --magnet              enable MAGNET filtering
  -mf, --magnet-filter <fa>  FASTA reference for filtering (mandatory with -mg)
  -mv, --magnet-verbose      MAGNET verbose
  -mt <0..1>                 similarity threshold (default: 0.9)
  -ml <1..44>                sensitivity level (default: 36)
  -mi, --magnet-invert       invert filter
  -mp <val>                  portion of acceptance (default: 1)
```

##### 18.2.3 Example (global + local profile)

```
./FALCON2 meta -v -F -l 47 -Z -y profile.com reads1.fq:reads2.fq VDB.fa
```

#### 18.3 Local detection and visualization

##### 18.3.1 `filter`: segment local interactions

```
./FALCON2 filter -h
```

```
Non-mandatory:
  -h  help       -F  force         -V  version       -v  verbose
  -s <size>      window size       -w <type>         window type
  -x <sampling>  window sampling   -sl <lb>          similarity lower bound
  -su <ub>       similarity upper  -dl <lb>          size lower bound
  -du <ub>       size upper        -t <thr>          threshold [0..2.0]
  -o <FILE>      output segmented filename

```

```
Mandatory:
  [FILE]        profile filename (from FALCON2 meta)
```

Example:

```
./FALCON2 filter -v -F -t 0.5 -o positions.pos profile.com
```

##### 18.3.2 `fvisual`: draw local/global similarity (SVG)

```
./FALCON2 fvisual -h
```

```
Non-mandatory:
  -h help  -F force  -V version  -v verbose
  -w <w>     square width               -s <is>   square inter-space
  -i <idxs>  color index start          -r <idxr> color index rotations
  -u <hue>   color hue                  -g <g>    color gamma
  -sl <lb>   similarity lower           -su <ub>  similarity upper
  -dl <lb>   size lower                 -du <ub>  size upper
  -e <sz>    enlarge painted regions
  -bg        best-of-group only
  -ss        hide global scale
  -sn        hide names
  -o <FILE>  output SVG filename

Mandatory:
  [FILE]     segmented filename (from FALCON2 filter)
```

Example:

```
./FALCON2 fvisual -v -F -o map.svg positions.pos
```

#### 18.4 Database inter-similarity

##### 18.4.1 `inter`: compute inter-similarity matrix

```
./FALCON2 inter -h
```

```
Non-mandatory:
  -h help   -V version  -v verbose  -s show levels
  -l <1..30>  compression level    -n <threads>
  -x <FILE>  similarity matrix     -o <FILE> labels

Mandatory:
  [FILE]:[FILE]:[...]   input FASTA files (':' to split)
```

Example:

```
./FALCON2 inter -v file1.fa:file2.fa:file3.fa
```

##### 18.4.2 `ivisual`: heatmap visualization

```
./FALCON2 ivisual -h
```

```
Non-mandatory:
  -h help  -V version  -v verbose  -w width  -a inter-space
  -s index start  -r index rotations  -u hue  -g gamma
  -l <FILE> labels  -x <FILE> heatmap (SVG)

Mandatory:
  [FILE]    input matrix (from FALCON2 inter)
```

Example:

```
./FALCON2 ivisual -F -l labels.txt -o heatmap.svg matrix.txt
```

### 19 Common use

A convenient end-to-end script:

```
#!/bin/bash
./FALCON2 meta   -v -n 4 -t 200 -F -Z -l 47 -y complexity.com $1 $2
./FALCON2 filter -v -F -t 0.5 -o positions.pos complexity.com
./FALCON2 fvisual -v -F -o draw.svg positions.pos
```

Make it executable and run:

```
chmod +x FALCON2-meta.sh
./FALCON2-meta.sh reads1.fastq:reads2.fastq VDB.fa
```

### 20 New features in FALCON2

#### 20.1 Model management

```
# Train + save
./FALCON2 meta -v -l 47 -S -M mymodel.bin -T reads.fq

# Load + infer
./FALCON2 meta -v -l 47 -L -M mymodel.bin reads.fq VDB.fa
```

Options: -S (save), -L (load), -M <file> (model file), -I (info), -T (train only).

#### 20.2 MAGNET integration

```
./FALCON2 meta -v -l 47 -mg -mf reference.fa -mt 0.9 -ml 36 reads.fq VDB.fa
```

Options: -mg (enable), -mf <fa> (filter reference, mandatory), -mv (verbose), -mt <0..1> (threshold, 0.9 default), -ml <1..44> (sensitivity, 36 default), -mi (invert), -mp <val> (portion).

### 21 Issues

Report problems at: GitHub Issues.

### 22 License

GPL v3. See LICENSE or GNU GPLv3.

© 2014–2025, IEETA, University of Aveiro.

### 23 Validation Results

Validation results for FALCON2 based on a battery of automated tests that verify correct operation across common and edge–case scenarios.

Each test asserts the program's exit status and logs full stdout/stderr for later inspection.

### 24 How to Run

```
# default (with valgrind, if available)
./test_falcon2.sh

# skip valgrind
NO_VALGRIND=1 ./test_falcon.sh
```

**Binary:** By default, the suite invokes ./FALCON2. Override with FALCON2=/path/to/FALCON2.
**Valgrind:** Enabled by default if present; disable with NO_VALGRIND=1.

#### 25 Artifacts and Logging

- **Run directory:** `./falcon2_logs/<YYYYMMDD_HHMMSS>/`
- **Per-test logs:** One log file per test case with the complete command, exit code, and program output.
- **Temporary data:** Synthetic FASTQ/FASTA test files (including empty, unreadable, malformed, and missing cases) are created under a temporary working directory and cleaned up automatically.

#### 26 Pass/Fail Semantics

- The script tracks total **PASS** and **FAIL** counts and exits non-zero if any test fails.
- Success/Failure is defined strictly by *exit status* comparisons to the expected code for each scenario.

```
./test_falcon2.sh
=========================================
  FALCON2 Comprehensive Test Suite
=========================================

Valgrind: DISABLED
Binary:   ./FALCON2
Version:   VERSION 3.2
Logs:     ./falcon_logs/20251025_213322
Root:     .

Generating synthetic test data...

[0] Early Exit Tests
  help (-h) ... PASS
  version (-V) ... PASS
  show levels (-s) ... PASS

[1] Training Mode (-T) - Synthetic Data
  train tiny reads ... PASS
  train small reads ... PASS
  train medium reads ... PASS
  train small.fq.gz ... PASS
  train medium.fq.gz ... PASS
  train empty file ... PASS
  train bad format ... PASS
  train missing file ... PASS
  train unreadable file ... PASS
  train no arguments ... PASS

[1b] Training Mode - Real Data
  train real reads.fq ... PASS
  train real reads.fq.gz ... PASS

[2] Model Save/Load/Info
  save model (-S -M) ... PASS
  save model different path ... PASS
  Model created: /tmp/tmp.XYJkgIg6Vc/trained_model.fcm (1409320719 bytes)
  load saved model ... PASS
  model info (-I) ... PASS
  load real model ... PASS
  real model info ... PASS
  load without -M flag ... PASS
  load missing model ... PASS
  save without -M path ... PASS

[3] Inference Mode - Synthetic Data
  infer tiny files ... PASS
  infer small files ... PASS
  infer medium files ... PASS
  infer small reads + medium db ... PASS
  infer small.fq.gz + small.fa.gz ... PASS
  infer mixed: fq + fa.gz ... PASS
  infer mixed: fq.gz + fa ... PASS
  infer only 1 arg (reads) ... PASS
  infer no arguments ... PASS
  infer missing reads ... PASS
  infer missing db ... PASS
```

```
  infer unreadable reads ... PASS
  infer invalid reads format ... PASS
  infer invalid db format ... PASS
  infer empty reads ... PASS
  infer empty db ... PASS
  infer both empty ... PASS

[3b] Inference Mode - Real Data
  infer reads.fq + VDB.fa ... PASS
  infer reads.fq + VDB2.fa ... PASS
  infer reads.fq.gz + VDB.fa.gz ... PASS
  infer reads.fq + VDB2.fa.gz ... PASS

[4] Magnet Filter Tests
  magnet basic filter ... PASS
  magnet verbose (-mv) ... PASS
  magnet with threshold ... PASS
  magnet with min length ... PASS
  magnet invert (-mi) ... PASS
  magnet max positions ... PASS
  magnet all params ... PASS
  magnet gz filter ... PASS
  magnet missing filter ... PASS
  magnet no -mf flag ... PASS
  magnet empty filter ... PASS

[5] Output File Tests
  output custom path (-x) ... PASS
  Output verified: /tmp/tmp.XYJkgIg6Vc/output1.csv (5 lines)
  output different path ... PASS
  output overwrite (with -F) ... PASS
  output readonly no -F ... PASS
  output with training ... PASS

[6] Thread Count (-n)
  threads n=1 ... PASS
  threads n=2 ... PASS
  threads n=8 ... PASS
  threads n=16 ... PASS
  threads n=32 ... PASS
  threads n=0 invalid ... PASS
  threads n=-1 invalid ... PASS

[7] Top Results (-t)
  top t=1 ... PASS
  top t=5 ... PASS
  top t=10 ... PASS
  top t=50 ... PASS
  top t=100 ... PASS

[8] Compression Level (-l)
  level l=20 ... PASS
  level l=30 ... PASS
  level l=36 ... PASS
  level l=40 ... PASS
  level l=47 ... PASS

[9] Gamma Parameter (-g)
  gamma g=0.000015 ... PASS
  gamma g=0.0001 ... PASS
  gamma g=0.001 ... PASS
  gamma g=0.01 ... PASS
  gamma g=0.1 ... PASS
  gamma g=0.5 ... PASS
  gamma g=0.9 ... PASS

[10] Collision Parameter (-c)
  collision c=1 ... PASS
  collision c=10 ... PASS
  collision c=50 ... PASS
  collision c=100 ... PASS

[15] Parameter Combinations
  combo: threads + top ... PASS
  combo: level + gamma ... PASS
  combo: all inference params ... PASS
```

```
  combo: training + save ... PASS
  combo: magnet + custom out ... PASS
  combo: local + profile ... PASS
  combo: sample + threads ... PASS
  combo: multi-file + magnet ... PASS

[16] Advanced Parameters
  local similarity (-Z) ... PASS
  profile output (-Z -y) ... PASS
  Profile created: /tmp/tmp.XYJkgIg6Vc/profile1.txt (6 lines)
  subsample p=1 (all) ... PASS
  subsample p=2 ... PASS
  subsample p=5 ... PASS
  subsample p=10 ... PASS
  subsample p=100 ... PASS
[17] Invalid Flags & Arguments
  unknown flag --invalid ... PASS
  typo flag -xyz ... PASS
  double dash alone -- ... PASS
  flag without value -n ... PASS

[13] Multi-File Input Tests
  multi reads: tiny:small ... PASS
  multi reads: small:medium ... PASS
  multi reads: 3 files ... PASS
  multi db: tiny:small ... PASS
  multi db: small:medium ... PASS
  multi db: 3 files ... PASS
  multi both: 2:2 ... PASS
  multi both: 3:2 ... PASS
  multi mixed: fq:fq.gz ... PASS
  multi mixed db: fa:fa.gz ... PASS
  multi all mixed ... PASS
  multi real: VDB:VDB2 ... PASS
  multi real: VDB:VDB.gz ... PASS
  multi real mixed: VDB.fa:VDB2.fa.gz ... PASS
  multi missing first ... PASS
  multi missing second ... PASS
  multi missing db first ... PASS
  multi empty in list ... PASS
  multi only colons ... PASS

[14] Edge Cases
  very small reads (1) ... PASS
  very small db (1) ... PASS
  both very small ... PASS
  thread count > cores ... PASS
  very high top value ... PASS

=========================================
  Test Summary
=========================================
Total:  132
Pass:   132
Fail:   0
Skip:   0
Logs:   ./falcon_logs/20251025_213322

Test Coverage:
  [0]  Early Exit Tests (help, version, show)
  [1]  Training Mode - Synthetic
  [1b] Training Mode - Real Data
  [2]  Model Operations (save, load, info)
  [3]  Inference Mode - Synthetic
  [3b] Inference Mode - Real Data
  [4]  Magnet Filter Integration
  [5]  Output File Handling
  [6]  Thread Count Parameter
  [7]  Top Results Parameter
  [8]  Compression Level Parameter
  [9]  Gamma Parameter
  [10] Collision Parameter
  [13] Multi-File Input (colon-separated)
  [14] Edge Cases
  [15] Parameter Combinations
  [16] Advanced Parameters (-Z, -y, -p)
```

```
  [17] Invalid Flags & Arguments

All tests passed!
==========================================
```
